# Robust parallel decision-making in neural circuits with nonlinear inhibition

**DOI:** 10.1101/231753

**Authors:** Birgit Kriener, Rishidev Chaudhuri, Ila R. Fiete

**Author notes:** B.K. and R.C. contributed equally to this work. Correspondence (I.R.F.).

## Abstract

Identifying the maximal element (**max, argmax**) in a set is a core computational element in inference, decision making, optimization, action selection, consensus, and foraging. Running sequentially through a list of *N* fluctuating items takes *N* log(*N*) time to accurately find the **max**, prohibitively slow for large *N*.

The power of computation in the brain is ascribed in part to its parallelism, yet it is theoretically unclear whether leaky and noisy neurons can perform a distributed computation that cuts the required time of a serial computation by a factor of *N*, a benchmark for parallel computation.

We show that conventional winner-take-all neural networks fail the parallelism benchmark and in the presence of noise altogether fail to produce a winner when *N* is large. We introduce the nWTA network, in which neurons are equipped with a second nonlinearity that prevents weakly active neurons from contributing inhibition. Without parameter fine-tuning or re-scaling as the number of options *N* varies, the nWTA network converges *N* times faster than the serial strategy at equal accuracy, saturating the parallelism benchmark. The nWTA network self-adjusts integration time with task difficulty to maintain fixed accuracy without parameter change. Finally, the circuit generically exhibits Hick’s law for decision speed. Our work establishes that distributed computation that saturates the parallelism benchmark is possible in networks of noisy, finite-memory neurons.

## Introduction

Finding the largest entry in a list of *N* quantities is an elemental and ubiquitous computation. It is invoked in a wide range of tasks including inference, optimization, decision making, action selection, consensus, and foraging^8;16;21;60^. In inference and decoding, finding the best-supported alternative involves identifying the largest likelihood (**max**), then finding the model corresponding to that likelihood (**argmax**); decision making, action selection and foraging involve determining and selecting the most desirable alternative (option, move, or food source, respectively) according to some metric, again requiring **max, argmax** operations.

Because **max, argmax** are building blocks in these myriad computations, it is important to characterize how long these operations take. The time-complexity of **max, argmax** or finding the largest number or its index in an unsorted list of *N* elements in an optimal serial procedure, as would be carried out on a computer, involves a sequential pass through the list, for a time linear in *N*. If each element is observed along with some noise, the computational complexity of solving the task with a desired accuracy increases to *N* log(*N*), as we will show. We will refer to these times as the *serial scaling* of max, argmax.

A major source of efficiency of computation in neural systems is hypothesized to be the potential for massive parallelism, with operations distributed across a circuit containing many neurons, thus trading computation time for space (i.e., number of neurons). However, it is not clear if a distributed computation spread across *N* neurons and the extraction of a final result from it can be acquired in a time that scales efficiently with *N*. In this paper, we will refer to a factor-*N* speed-up relative to the optimal serial strategy as the *parallelism benchmark*.

There are two distinct regimes in which it is interesting to consider how (fast) the brain computes **max, argmax**: The first is finding the most active neuron (or neuron pool) across thousands of neurons (pools). An example of this large-*N* **max, argmax** computation is the dynamics that lead to the sparsification of Kenyon cell activity within the fly mushroom bodies^60^, and potentially to the sparse activation of hippocampal place cells^1;17;59^. It is possible that many more areas with strong recurrent inhibition and gap-junction coupled interneurons display similar dynamics, including the vertebrate olfactory bulb^49;58^ and basal ganglia^47;53^. The second regime is encountered in explicit behavioral decision making across a small number of externally presented alternatives. We seek an understanding of how neurons compute **max, argmax** across these disparate regimes.

Because of the importance of **max, argmax** operations, they are well-studied in neuroscience in the guise of *winner-take-all* (WTA) neural circuit models and phenomenological *accumulate-to-bound* (AB) models. A WTA network consists of N neurons (pools) driven by external inputs, each amplifying its own state and interacting competitively through global inhibition. Self-amplification and lateral inhibition can produce a final state in which only the neuron with the largest integrated input (**argmax**) remains active, while the rest are silenced^10;22;39^. If the activation of the winner is proportional to the size of its input^69^, the network also solves the **max** problem. The final state of the network serves as the completed output of the computation.

By contrast, AB models^6;31;43;51;64^ consist of individual integrators that sum their inputs. Unlike WTA models, AB models require a separate downstream readout, which must decide which integrator has largest value, typically done by applying a threshold. Thus they do not, by themselves, output an answer to **max, argmax**. AB models are phenomenological rather than neural, and unlike WTA models typically do not incorporate the constraints of leaky, interacting neurons. For all these reasons, our focus is on WTA networks.

Despite the elemental nature of WTA computation in neural circuits, the time-complexity of WTA — how long it takes a system to compute **argmax** and **max** from a set of *N* inputs, as a function of *N* — is not well-characterized (see Discussion). On the one hand, one might expect that the parallel architecture of neural networks could speed up the computation. On the other hand, neural elements are hardly ideal parallel processors or integrators: they are leaky, and their nonlinear thresholds combined with noise could discard relevant information. Thus, it is unclear whether parallel processing with neurons can manage the time-for-space tradeoff efficiently, reducing temporal complexity by an amount in proportion to the increase in spatial complexity and thus achieving the parallelism benchmark.

We show that conventional winner-take-all circuit models in the presence of noise either fail to approach the parallelism benchmark or altogether fail to produce a winner when *N* is large. We propose the nWTA network, in which neurons are equipped with a second nonlinearity so that weakly active neurons cannot contribute inhibition to the circuit. The nWTA network can both match the accuracy of the serial strategy, and do so *N* times faster, saturating the parallelism benchmark. Unlike the conventional models, this performance requires no parameter fine-tuning. Moreover, the nWTA network self-adjusts (without any parameter change) its integration time as log(*N*) with the number (*N*) of noisy options, matching both Hick’s law of behavioral decision making^25^ and a normative scaling of fixed-accuracy decision making with noisy options.

In total our results suggest that it is at least theoretically possible for neural circuits to perform and exploit truly parallel computation.

## Results

### The problem

Consider the problem of finding the largest element in a set of *N* scalar-valued options of magnitudes *b*_1_ > *b*_2_ ≥ *b*_3_… ≥ *b_N_*, numbered in descending order of strength. In the deterministic case, the options are presented without noise. In the noisy case, the options fluctuate over time, 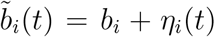, where *b_i_* is the fixed mean and *η_i_*(*t*) are zero-mean fluctuations (see Methods), and the task is to identify the item with the largest fixed mean. We assume that the top two options are separated by an N-independent gap of size Δ, so that *b*_1_ = *b*_2_ + Δ. For simplicity, we also assume that the remaining inputs are equal to each other (b = (*b* + Δ, *b*,…, *b*)^τ^ and *b*, Δ > 0). For a discussion of more general distributions of the inputs (including a fully uniform distribution, i.e., all *b*_i_ ~ U[0,1] where the top gap shrinks with *N*), see SI S2.3, S2.4.

### Optimal serial strategy, parallelism benchmark, and Hick’s Law

We first consider theoretical bounds on the decision time for **max, argmax** and compare these bounds with human behavior. Then, we will consider how neural circuits perform relative to the bounds.

Consider the time complexity of a serial procedure, as would be carried out on a computer. As described earlier, the time taken to choose the largest of a set of N non-noisy options presented serially takes a time 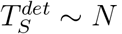. This is the *deterministic serial scaling* of **max, argmax**.

In the noisy case, obtaining a correct answer involves collecting information for long enough to estimate the input means, then performing a **max** operation on the estimates. Any strategy with a finite decision time will have a non-zero error probability, and we expect a *speed-accuracy tradeoff* that involves setting an acceptable error probability and finding the shortest time required to reach this accuracy (one may also make a maximally accurate decision within a set time *T*; see S1.2).

Consider the (*N* – 1) summed differences 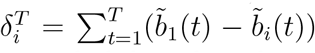 between the highest-mean input and each of the others. The probability that the wrong element is selected after averaging for time *T* is bounded by the sum of the probabilities that any individual *δ_i_* is less than zero. The quantities *δ_i_* concentrate around the true gaps (Δ*_i_* = *b*_1_ – *b_i_*), with the probability of error-inducing fluctuations decaying as 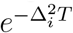 across a wide set of input distributions, Figure 1a and S1.1. The waiting time *T* ~ log(*N*)/Δ^2^ depresses individual error probabilities so they scale as 1/*N*, keeping the total error probability constant with *N*, Figure 1b and S1.1. Thus, the time for a serial strategy to achieve a constant decision accuracy is bounded by 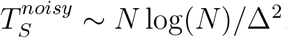. This is the *noisy serial scaling* of **max, argmax**.

**Figure 1:**
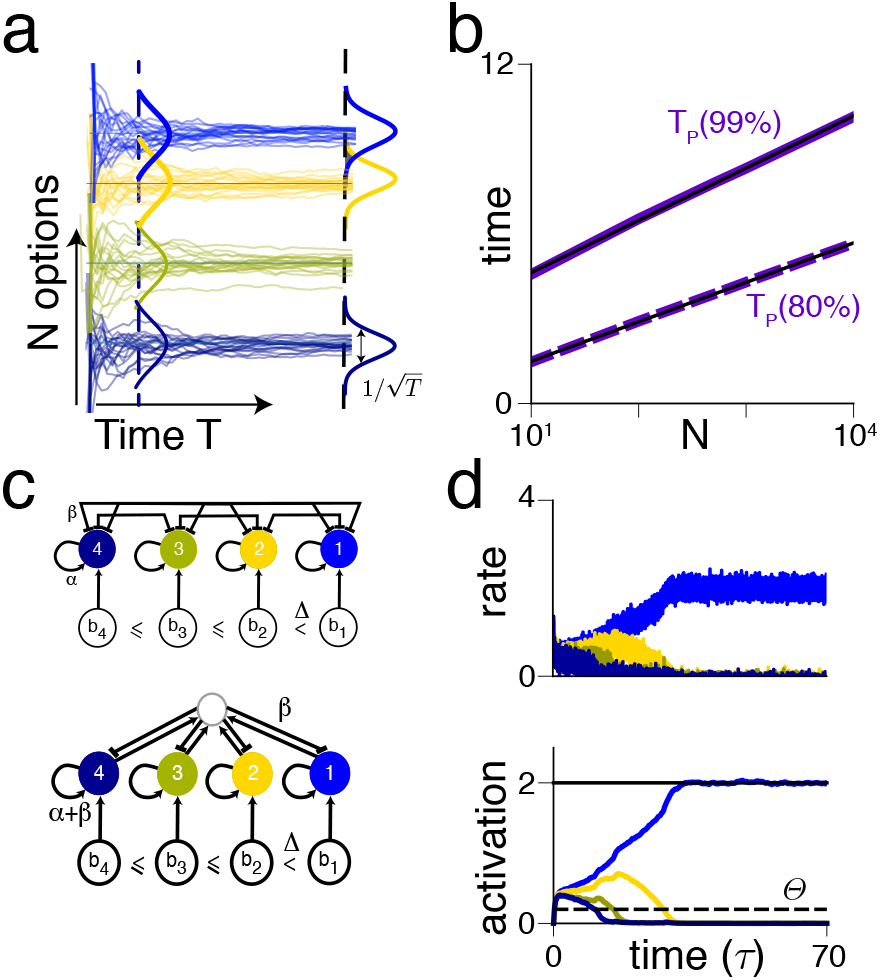
Parallelism benchmarks and setup of the WTA circuit. (a) Schematic illustration of how log(*N*) integration time is sufficient to maintain fixed accuracy. Thin lines: samples from each option (*N* options are in different colors). Thick curves: sample histograms (distributions) after integrating for time *T*; the standard deviation shrinks as 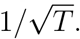. (b) The parallelism benchmark requires log(*N*) integration time to maintain fixed accuracy with noisy inputs. Plot shows the integration time to fixed accuracy (dashed and solid: 80%, 99% accuracy respectively) in (non-neural) simulations with Gaussian input fluctuations. Thin black lines: logarithmic fits. Matching analytical results using concentration inequalities are in S1.1. (c) WTA network architectures. Top: Neurons (or pools of neurons), ordered by the size of their external input (with gap Δ ≡ *b*_1_ – *b*_2_ between the top two inputs), inhibit all others and excite themselves. Bottom: Mathematically equivalent network with a global inhibitory neuron (pool): all neurons (pools) excite the inhibitory neuron, which inhibits them. This requires only ~ *N* synapses, compared to *N*^2^ for the mutual inhibition circut (top). (d) Neural firing rates and synaptic activations (coloring as in (c)). Convergence time *T*_WTA_ is the time taken for the firing rate of the winner neuron to reach its expected asymptotic activity range (black line, see Methods). Dashed line: threshold for a neuron to contribute inhibition in the nonlinear inhibition model (nWTA, see Methods).

A strategy that can process options in parallel (rather than considering each in turn) should achieve a factor of *N* speedup over the serial strategies, achieving times 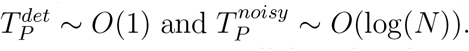 We will refer to these idealized parallel computation times as *parallelism benchmarks*.

This theoretical benchmark on parallel **max** with noisy options matches the empirical Hick’s law^6;25;33;43;44;63–65^, an influential result in the psychophysics of human decision making used as far afield as commercial marketing and design to improve the presentation of choices^34^. Hick’s law states that the time to reach an accurate decision increases with the number of alternatives, *N*, as log(*N* + 1). Our derived parallelism benchmark therefore suggests that the phenomenology of Hick’s law might reflect a general process of efficient decision-making when options are noisy.

The origin of log(*N*) scaling in the need to average away noise in our result is in contrast to existing explanations of Hick’s Law, which attribute the scaling to the number of progressive binary classification steps needed to winnow *N* (deterministic) options down to one, or equivalently to the number of bits required to specify one out of *N* options^25;50^. These alternative explanations would predict Hick’s law-like log(N)-scaling also for deterministic options, while our theory predicts a crossover from logarithmic scaling of decision time with number of options to a time that is independent of the number of options when the task is sufficiently noiseless or noise is reduced through practice. This crossover prediction is consistent with some studies^50;63^, and could provide a good test of whether Hick’s law is fundamentally due to noise in the alternatives during decision making.

The natural question, which we address in the rest of what follows, is what type of neural circuit can perform such efficient parallel decision making between noisy options?

### Neural decision-making circuits

The parallelism benchmarks derived above are idealizations without a neural implementation. The benchmarks require perfect integration of inputs without leak, and assume no loss of information from internal noise and nonlinear processing, as neurons might contribute. They are also not self-terminating: they require an external observer to apply a threshold to terminate the operation and select the option on top at that time as the winner.

The natural way to model a self-terminating **max, argmax** computation in the brain is through the *winner-take-all* (WTA) circuit, with a variety of circuits in the brain exhibiting WTA-like architectures. Consider *N* neurons (or neuron pools) interacting through self-excitation (strength *α*) and mutual inhibition (strength *β*), Figure 1c. The neural states are described by synaptic activations *x_i_*(*t*) or firing rates *r_i_*(*t*), i ∈ {1,…*N*}:

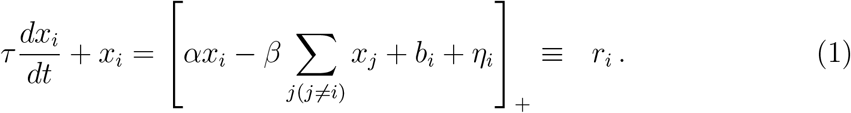

Here, [·]_+_ = max[0, ·] is a rectification nonlinearity, and *b_i_* and *η_i_* are the mean and fluctuations in the inputs, as described previously (the fluctuations may arise from input noise, stochastic neural activity, or both). For appropriate parameter values, and in the absence of noise, the network exhibits winner-take-all dynamics with a unique winner^69^, Figure 1d (and S2.1). These dynamics can be understood as movement downhill on an energy landscape, which drives the network to one of *N* possible stable states, each corresponding to solo activation of a different neuron (Figure S1).

We wish to understand how WTA dynamics behave as a function of the number of competing alternatives *N*, to arrive at an accurate single estimate of a winner (*T*_WTA_).

### Conventional WTA networks do not show efficient parallel decision-making

#### Weak inhibition: Accurate but slow WTA

Total inhibition in Equation (1) grows as *βN*. A reasonable possibility is to scale inhibitory strengths as *β = β*_0_/*N*, where *β*_0_ is some constant. We call this “weak” inhibition (Figure 2a).

**Figure 2:**
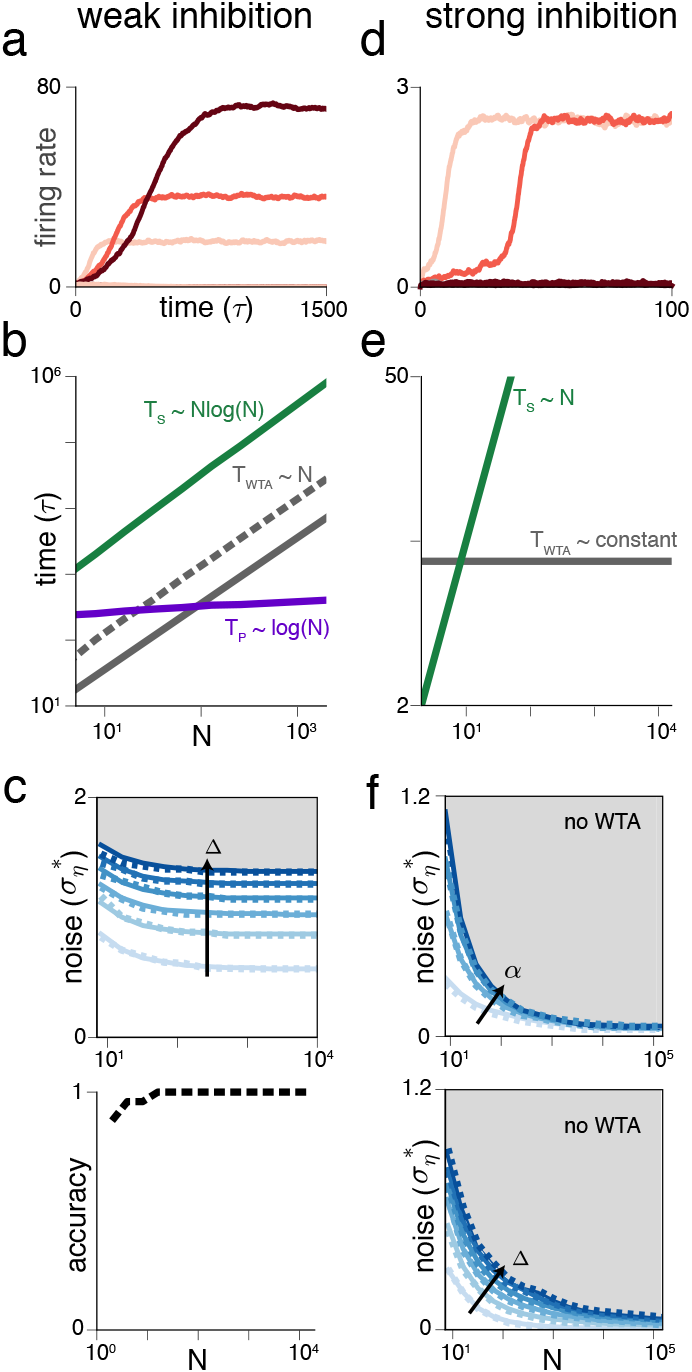
Conventional WTA networks fail the parallelism benchmark. (a)-(c) Results for weak inhibition. (a) Activity dynamics of the most-active neuron for networks of size N = 10, 20, 40 (light to dark red), respectively, with noisy input. [*α* = 0.6; *β* = 1; Δ = 0.1; *σ_η_* = 0.35]. The asymptotic activity level grows with *N*. (b) WTA simulation results on decision time without (solid gray) and with (dashed gray) noise. Also shown: noisy serial strategy (green) and noisy parallelism benchmark (purple). Error bars are smaller than the linewidth. Deterministic serial strategy (i.e., O(*N*)) not shown for simplicity. (c) Top: Critical noise amplitude versus *N*: WTA dynamics exists below a given curve and fails above it (dashed: numerical simulation; solid: analytical). Darker curves correspond to a larger gap between top two inputs. Bottom: Average accuracy as function of *N*. (d)-(f) show results for strong inhibition. (d) As in (a) but for strong inhibition. WTA breaks down rapidly (note absence of WTA for *N*=40, dark curve). (e) WTA decision time in the absence of noise (gray) along with the deterministic serial strategy (green). (f) As in (c), showing critical noise amplitude versus *N*. Darker curves: stronger selfexcitation (top; α = {0.1,0.6,0.9}; Δ = 0.1) or a widening gap (bottom; Δ = {0.01,0.06,…, 0.26}; *α* = 0.6). [*β* = 1 in all curves.]

Self-excitation (*α*) must be set to both maintain stability and assure a WTA state, requiring 1 – *β*_0_/*N* < *α* < 1^69^, S2.1. The two-sided constraint is a *fine-tuning* condition: for large N, the allowed range of self-excitation is very small and shrinks towards zero.

In the deterministic case the network always converges to the correct solution where the neuron with input *b* + Δ is the winner. The equations can be analytically solved to obtain the convergence time (S2.2):

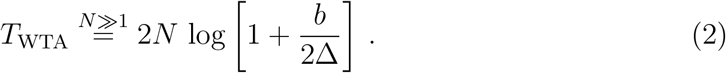

The linear growth of *T*_WTA_ is the same as the serial strategy, Figure 2b, and offers no parallel speedup.

Above a certain critical value of noise (predicted analytically, S2.5), the weak inhibition network fails to exhibit WTA dynamics, Figure 2c top panel. This failure is to be distinguished from an error: The network does not select the wrong winner, it simply fails to arrive at a winner and multiple neurons remain active. Below the critical noise threshold, WTA dynamics persists even for *N* → ∞, Figure 2c top panel. The critical noise threshold is substantially larger than Δ, and the network exhibits WTA even when Δ = 0 (lightest line in Figure 2c), selecting a random neuron as the winner. The decision time with fluctuating inputs (Figure 2b) grows linearly as for deterministic inputs (see S2.2 for an explanation of the linear scaling). Though this time-scaling is much slower than the parallelism benchmark, it asymptotically (large *N*) exhibits perfect computation accuracy, Figure 2c bottom panel (the *N* above which perfect accuracy is obtained depends on Δ and *σ_η_*).

In summary, the existence of WTA in a weak-inhibition circuit requires exquisite fine-tuning of excitation and cannot be adjusted to provide a speed-accuracy tradeoff. The network favors accuracy over speed, always achieving perfect accuracy for large enough *N*, but the convergence time grows as ~ *N*, a modest speed-up of log(*N*) relative to 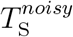 that does not approach the factor of *N* speed-up of an efficient parallel strategy.

#### Strong inhibition: parallel speedup without noise, but WTA breakdown with noise

An alternate choice is to hold *β* fixed as *N* is varied. We call this “strong” inhibition (Figure 2d). The total inhibition in the circuit then grows with *N*. A unique WTA solution exists for any choice of *α* in the *N*-independent interval (1 – *β*, 1] (no fine-tuning required, unlike weak inhibition).

For noiseless inputs, we analytically obtain (S2.2):

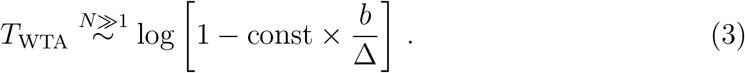

As with the weak inhibition case, *T*_WTA_ depends on Δ. Notably, however, *T*_WTA_ is independent of *N*, Figure 2e, solid black, matching the parallelism benchmark.

In the noisy case, for a given *N*, the network can perform WTA if the noise amplitude is smaller than a threshold that depends on Δ. If *N* grows (while holding Δ and noise amplitude fixed), however, the network entirely fails to reach a WTA state, Figure 2f. Unlike with weak inhibition, strong inhibition networks appear to fail to asymptotically exhibit WTA behavior for any non-zero noise level.

We can understand the failure as follows. Unbiased (zero-mean) noise in the inputs, when thresholded, produces a biased effect: neurons receiving below-zero mean input will nevertheless exhibit positive mean activity because of input fluctuations. Thus, even neurons with input smaller than their deterministic thresholds continue to inhibit others. Total inhibition remains ~ *N* over time and, for sufficiently large *N*, prevents any neuron from breaking away from the rest to become a winner.

As with weak inhibition, we can analytically predict the breakdown of strong-inhibition WTA, S2.5. The resulting predictions can be used to determine the feasibility of WTA computation in large networks in the presence of noise.

Thus, although networks with strong inhibition can meet the parallelism benchmark when the inputs are deterministic, they are not capable of finding a winner in large networks with even slightly noisy inputs (and when they do perform WTA for sufficiently small N, the scaling is suboptimal; data not shown). These results are pessimistic and raise the question of whether neural networks can ever implement parallel computation that is efficient, fully trading serial time for space.

### The nWTA network: fast, robust WTA with noisy inputs and an inhibitory threshold

We introduce a new model motivated by the successes and failings of the existing models. A network with weak inhibition and fine-tuning is accurate but too slow because inhibition is too weak to drive a rapid separation between winner and losers. A network with strong inhibition achieves WTA with a full parallelism speed-up for deterministic inputs, but entirely fails to perform WTA for large *N*, because the nearly-losing neurons prevent any neuron from breaking away. (Simply increasing the activation threshold for all neurons does not fix the failure of WTA, Figure S3e). In addition, the residual noise-driven inhibition decreases the average asymptotic activity of the near-winner so that the true value of **max** will be underestimated.

Consider a circuit with strong inhibition, in which individual neurons can only contribute inhibition when their activations exceed a threshold *θ* (see Discussion for biological candidates):

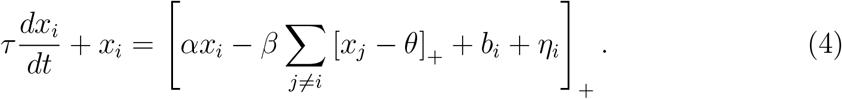

We set 0 < *θ* < *b*_1_/(1 – *α*) (no fine tuning) and self-excitation in the range 1 – *β* < *α* < 1.

In this *nonlinear-inhibition WTA* (nWTA) network, if the ith neuron wins, its expected asymptotic state is *b_i_*/(1 – *α*), so the direct proportionality to **max** is recovered. Every neuron contributes an inhibition of strength ~ 1 when highly active (> *θ*), ensuring robust early competition. However, the inhibitory threshold will cause neurons with decreasing activations to effectively drop out of the circuit, with the hope that the top neuron could take off unimpededly later in the dynamics.

When presented with deterministic inputs, the nWTA network matches the conventional strong inhibition network and the parallelism benchmark by converging in a time independent of N, Figure 3a.

**Figure 3:**
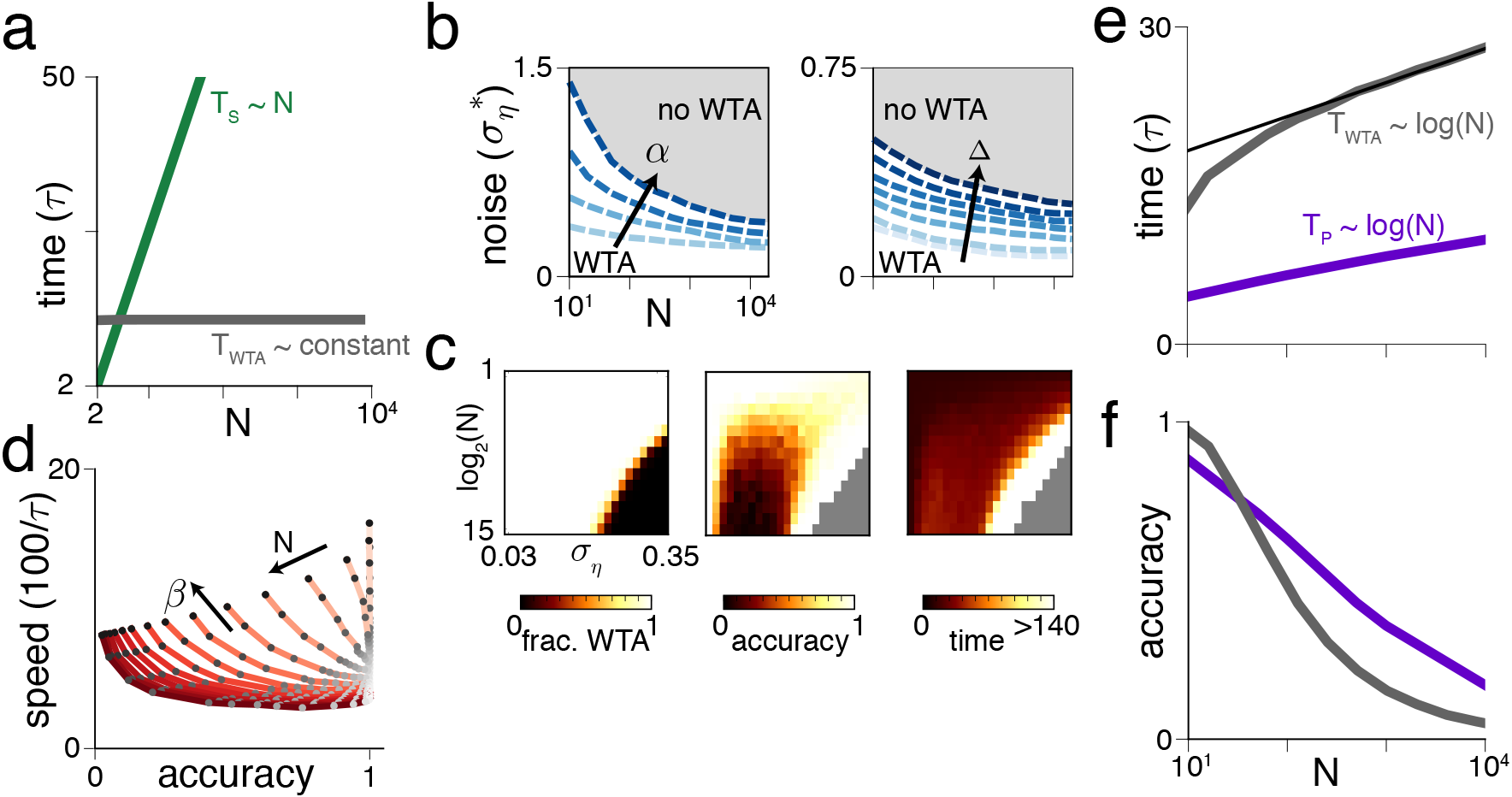
Nonlinear inhibition WTA (nWTA) is robust to noise and achieves the parallelism benchmark. (a) nWTA decision time without noise (gray) and the deterministic serial strategy (green). (b) Critical noise amplitude 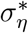 as a function of *N* for varying *α* = {0.5,0.7,0.9,1.1}, Δ = 0.075 (left) and Δ = {0, 0.0125, 0.025, 0.05, 0.075, 0.1, 0.15}, *α* = 0.5, *β* = 0.51 (right). Below each curve, WTA behavior exists, while above it does not. (c) Heatmaps showing fraction of trials with a WTA solution (single winner; left), accuracy of the WTA solution (middle) and convergence time *T*_WTA_/*τ* (right) as function of network size *N* and noise amplitude *σ_η_*. Δ = 0.05, *α* = 0.5, *β* = 0.6. (d) Speed-accuracy curves for Δ = 0.075, *σ_η_* = 0.12 for varying *N* = {2^3^, 2^4^,…, 2^15^} (light to dark red) and *β* = {0.51, 0.52,…, 0.6, 0.65, 0.7,0.8, 0.9} (light to dark gray circles). Only trials that produced a WTA solution were included. Curves are non-monotonic, so that certain parameters are strictly better than others for both speed and accuracy. (e) N-scaling of decision time to achieve fixed accuracy of 0.99 for nWTA (gray) and the parallelism benchmark (purple). Thin black line shows logarithmic fit. [Δ = 0.075; *τ_η_* = 0.05*τ*; *θ* = 0.2; *σ_η_* = 0.12; see S2.6, Figure S4 for similar results with different parameters and noise levels.] (f) *N*-scaling of accuracy at fixed decision time. Gray: nWTA for integration time 12.5*τ*; purple: parallelism benchmark for integration time 2*τ*.

Moreover, and unlike the conventional WTA network, diminishing inhibition in the nWTA circuit over time permits the leading neuron to break away and win even in the presence of noise. The losing neurons continue to receive inhibitory drive from the remaining highly active neuron(s), becoming less active over time. The network exhibits WTA behavior well into the noisy regime even with asymptotically many neurons, Figure 3b,c, without fine-tuning (see S2.5, Fig S3f for more discussion).

The network can also exhibit a broad tradeoff between speed and accuracy. Starting at high accuracy and holding noise fixed, the accuracy of computation can be decreased, and speed increased, by increasing *β* (Figure 3d, for fixed *α*; darker gray circles along a curve correspond to increasing *β*) or *α* (Figure S4a-f). The overall integration time is generally set by the combination of *α* and *β*, with high accuracy and low speed achieved as *α* + *β* approaches 1. When *α* + *β* is increased away from 1, the overall trend is that speed increases and accuracy decreases. Note, however, that the nWTA network exhibits an interesting non-monotonic dependence of accuracy on both noise level and network size (as can be seen in the heatmap of Figure 3c). This improvement in performance at some higher noise-level is a form of stochastic resonance^42^ (see S2.6 and 2.7 for discussion).

Conveniently, a top-down neuromodulatory or synaptic drive can regulate where the network lies on the speed-accuracy curves, with many mechanistic knobs for control, including synaptic gain control of all (excitatory and inhibitory) synapses together (resulting in covariation of *α,β*), neural gain control of principal cells (also effective covariation of *α,β*), or a threshold control of inhibitory cells (effective modulation of *β*).

For fixed noise at each input or WTA circuit neuron, the decision time for the network to reach a fixed accuracy scales as *T*_WTA_ ~ log(*N*) (Figure 3e; also see Figures S4e,k). Compared to the serial time-complexity of 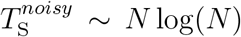 for fixed accuracy with noisy inputs, the nWTA network therefore achieves a fully efficient tradeoff of space for time, matching the parallelism benchmark of 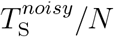.

Not only does the *scaling* of decision time with *N* in the nWTA network match the functional form of the parallelism benchmark, the prefactor is not far from unity: it takes only a factor of ~ 3 more time steps (in units of the biophysical time-constant of single neurons) to converge than the number of discrete time-steps of the parallelism benchmark (Figure 3e, black vs purple curves).

Similarly, if we compare accuracy at fixed *T*_WTA_ in Figure 3f (here at *T*_WTA_ = 12.5*τ*) to the parallelism benchmark at some fixed *T_s_/N = kT*_WTA_ (here, *k* = 0.16), we see that accuracy at large *N* is similar to that of the parallelism benchmark, again given a near-unity prefactor (Figure 3f). Further, as we show in the following sections, for small *N* even the prefactor can be almost optimal.

In summary, the nWTA network can perform the **max, argmax** operations on noisy inputs with comparable accuracy as the optimal serial strategy, but with a full factor-*N* parallelism speed-up, even though the constituent neurons are leaky. It does so with network-level integration and competition, but does not require fine-tuning of network parameters.

### Self-adjusting dynamics: high accuracy and Hick’s law without parameter tuning

The results above on efficient parallel decision-making by nWTA across a wide range of *N* assumed that network parameters can be changed to optimize the decision time for the given value of noise or *N*. As previously mentioned, there are many biologically-plausible mechanistic ways to change effective excitation, inhibition and thresholds in the network. Nevertheless, the assumption that the network can retune parameters for each new set of inputs may not hold in all cases. The re-tuning assumption may be unrealistic for psychophysical decision-making among small numbers of alternatives, where the number and statistics (noisiness) of inputs are controlled by the external world and may change from trial to trial and without prior warning. We thus investigate the performance of nWTA when parameters do not change with *N*.

Remarkably, nWTA networks can self-adjust to maintain high accuracy as *N* increases up into the thousands, with an appropriate fixed initial parameter choice, Figure 4a. The network maintains high accuracy as *N* increases by automatically extending its decision time, Figure 4b. The increase in *T*_WTA_ results from recurrent inhibition, which slows convergence based on the number of active options. The automatic increase in *T*_WTA_ is logarithmic in *N*, matching the parallelism benchmark and reproducing Hick’s Law, Figure 4b.

**Figure 4:**
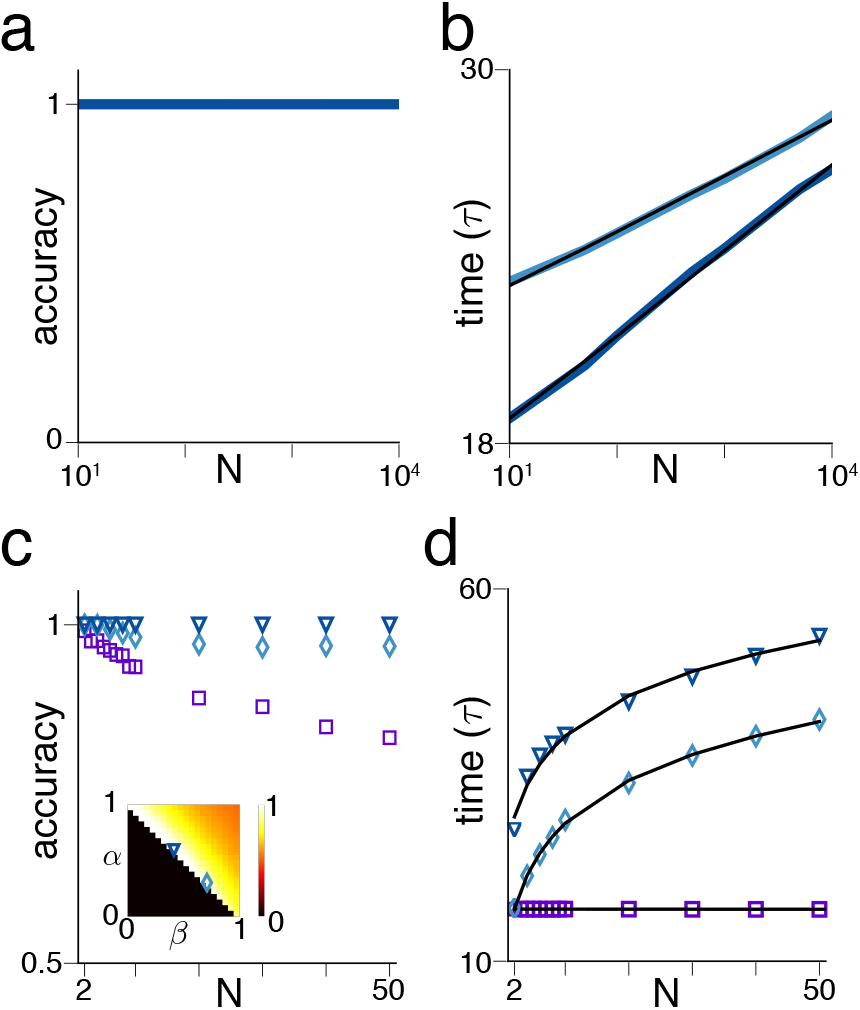
nWTA network self-adjusts to maintain accuracy and exhibits Hick’s law without any parameter change. (a) Accuracy for fixed parameter values, as a function of N. Dark and light blue show (*α, β*)=(0.5, 0.51), (0.6, 0.41) respectively (one trace not visible). Input parameters as in Figure 3d-f. (b) Decision time for the parameter values shown in (a), along with logarithmic fits (thin black). (c) Accuracy for fixed parameter values for small *N*. Blue symbols show nWTA network (light, dark show (*α, β*) =(0.3, 0.71), (0.6, 0.41) respectively) while purple squares show AB model with perfect integration. Δ = 0.05, *σ* = 0.2. Inset shows selected parameter values as well as average accuracy for all parameter combinations (averaged across *N* = 2 – 10). (d) Decision time for nWTA networks and AB model shown in (c). Thin black lines show logarithmic fits for nWTA and constant fit for AB.

For the small-*N* case relevant for psychophysical decision-making, the ability to maintain high accuracy without parameter re-tuning is highly robust: nWTA networks show high accuracy across a range of (fixed) possible parameter settings, Figure 4c (blue symbols). Moreover, for every parameter combination we examined, the increase of *T*_WTA_ with *N* was better described as logarithmic than linear, e.g. Figure 4d (blue symbols, and Figure S6j for full plane), regardless of whether high accuracy was maintained. Thus, Hick’s law is a generic dynamical consequence of decision-making by self-terminating WTA networks with nonlinear recurrent inhibition.

The observed increase in decision time with *N* can be reproduced by accumulate-to-bound (AB) models of decision-making, in which the evidence for each option is integrated by either perfect or leaky accumulators. These models integrate noisy input to an externally-imposed threshold that must be hand-designed to scale as log(*N*) in order to reproduce Hick’s law^43;64^. However, when the parameters of these models are kept fixed with *N*, the behavior is qualitatively different from WTA networks: the decision time remains fixed with *N* (instead of increasing logarithmically), and accuracy decreases, Figure 4c,d (purple squares). Thus, if *N* and the appropriate threshold are not known ahead of time so that the model threshold cannot be retuned, AB models show behavior very different from Hick’s Law. By contrast, the nWTA model maintains high accuracy and shows Hick’s Law behavior while in absolute terms taking only slightly longer than perfectly integrating AB models to reach a matching accuracy, Figure 4c,d.

Thus we have found both a computational (efficient normative computation that saturates the parallelism benchmark) and dynamical explanation (self-adjustment of decision time with number of options without parameter change) of Hick’s law.

### WTA as a minimal, flexible model for multi-alternative forced-choice decision making

Despite its simplicity, the nWTA model can not only maintain performance comparable to idealized, leakless AB models as shown above and elaborated next, but can also reproduce a number of behavioral and physiological observations from multi-AFC studies. These include faster decision times on correct compared to error trials, the flexibility to preferentially weight early evidence, the existence of step- and ramp-like responses of individual neurons, a higher pre-decision activity under speed pressure, convergence to the same activity level at decision time even though *N* is varied, and partial self-adjustment to the statistics of the input. Throughout this section, the parameters used are comparable to the parameters used to achieve high performance in the large-N regime, suggesting that the same basic circuit can be reused across a variety of scales.

The nWTA circuit can achieve a broad range of speeds and accuracies, with a corresponding range of reward rates (i.e., product of speed and accuracy, a key quantity for psychophysics experiments; throughout we assume an additional non-decision time *T*_0_ = 300 ms, see S3), by varying self-excitation and recurrent inhibition, Figure 5a,b. The reward rates achieved by the self-adjusting dynamics of nWTA networks when parameters are held fixed as *N* is varied are comparable to when they are separately optimized for each *N*, Figure 5a (dark versus light symbols). The self-adjustment process is thus near-optimal, at least over the range of number of options considered here (2 ≤ *N* ≤ 10).

**Figure 5:**
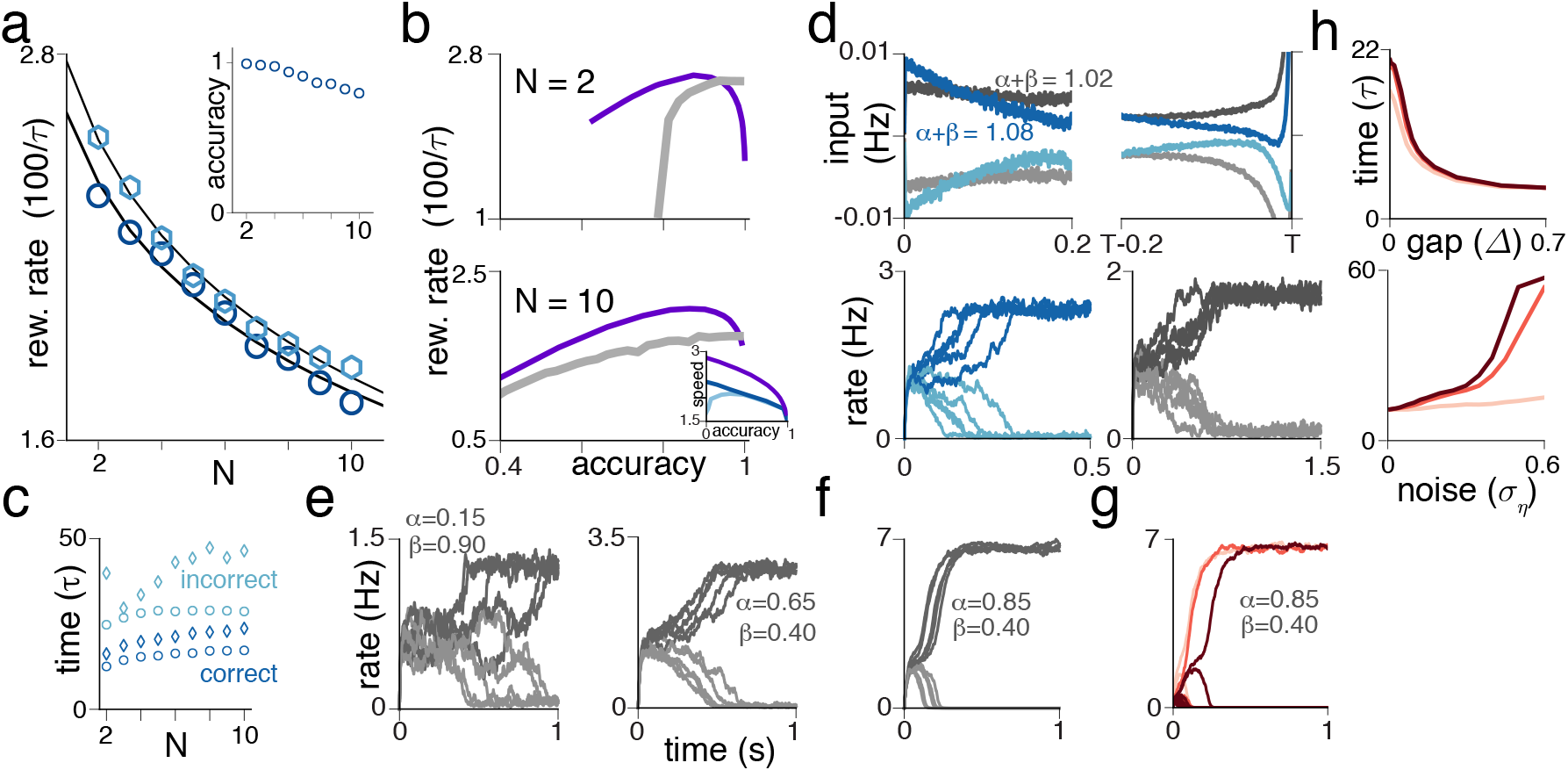
Self-terminating WTA dynamics as a minimal-parameter, neural model of multi-AFC decision-making. (a) Reward rate for the fixed parameters chosen to maximize reward rate (dark blue circles, *α* = 0.41, *β* = 0.7) along with reward rate when parameters are individually optimized for each *N* (light blue). Thin black lines show fit to (*T*_0_ + log(*N* + 1))^-1^. Inset shows accuracy for fixed parameters chosen to maximize reward rate. Accuracy at highest reward rate is high but not constant, reflecting a speed-accuracy tradeoff. (b) Reward rate vs. accuracy for *N* = {2,10}. Thick gray: nWTA (using best *α, β* for given accuracy); purple: AB model (note that AB model takes time 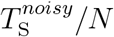 and is thus the parallelism benchmark). Inset for *N* =10 panel shows speed-accuracy curve (dark blue: *α* fixed, *β* varied; light blue: *β* fixed, *α* varied; *T*_0_ = 300 ms). (c) Convergence time for correct vs. incorrect trials (circles, diamonds show (*α,β*) = (0.41, 0.7), (0.46, 0.6) respectively).(d) Top: average input for winning (dark gray; blue) and losing (light gray; blue) neurons in an nWTA network performing a 2-AFC task with zero gap for two parameter settings (gray versus blue). Bottom: example firing rates from trials used for the averages in the top panel. (e) Example firing rate trajectories for nWTA networks with different (*α,β*) but with *α + β* (i.e.,integration time-constant) held fixed. Left: lower self-excitation and higher inhibition; right: vice versa. (f) Example firing rate trajectories for network with same inhibition as (e) (right panel) but stronger self-excitation. Trials are faster, but final activation of winner is higher. (g) Firing rate trajectories for *N* = 2, 6,10 (light to dark red) with fixed parameters. Decision threshold remains constant. (h) Top: Decision time for fixed parameters as a function of gap between largest and second largest input (related to task difficulty), showing that network self-adjusts to integrate longer for difficult tasks. Colors as in (g). Bottom: As in top panel but for noise in input. Note that in the small-N regime, conventional WTA models show similar performance to nWTA for many of these results, though less robustly; see SI Figure S6 and S3.

Moreover, the reward rate achieved by nWTA networks, both when parameters are held fixed, or when they are individually optimized for each *N* is competitive with the non-leaky benchmark parallel AB models (Figure 5b, cf. gray for network and purple for AB model, see also Figure S6b). The nWTA network even outperforms parallel AB in the high-accuracy regime at small *N* (see^43^ for a similar result; see also Figure S6b and S1.4 for discussion of a different non-parallel AB benchmark that performs better than parallel AB and nWTA at small *N* through competition, but is biologically unrealistic and no better than parallel AB for larger *N*).

When parameters and *N* are held fixed, correct WTA trials terminate faster than wrong ones (Figure 5c), as observed in other attractor-based decision models^19;68^ and consistent with psychophysics^38;54^. By contrast, AB models do not produce the more-accurate-is-faster result^41;51;61^ without additional modifications ^52^.

The WTA condition *α + β* > 1 corresponds to unstable network dynamics. Thus, the network is generically impulsive, more strongly weighting early inputs relative to late ones ^27;30;67^, as seen in the decision-triggered average input curves of Figure 5d (top row; blue curves). It is difficult to achieve leaky integration with a self-terminating decision process without further modification of the nWTA circuit.

However, the network can achieve near-uniform integration when tuned, with *α + β* set close to 1, Figure 5d (top row, gray curves). To obtain a uniform weighting of evidence over the 1-2 second integration time-windows tested in existing experiments^7;29;57^, the tuning of *α + β* to 1 need not be finer than ~ 2%, even if the biophysical time-constant of single neurons or synapses is as short as 20 – 50 ms (see Figure S1). The tuning required to arrive at this or a more finely specified parameter setting is likely achieved through plasticity during task training. AB models, by contrast, do not naturally display impulsive dynamics, instead requiring the addition of another dynamical process, such as an “urgency signal” or collapsing decision thresholds over time during the trial, to reproduce impulsive behavior^9;11^.

During integration, neurons vary enough from trial to trial for fixed parameter settings and across parameter settings to look variously more step-like or ramp-like (compare curves within and across Figure 5d-f). Different choices of *α,β* modulate the neural response curves, shifting them from more ramp-like to more step-like even as the network integration time (*τ*/(1 – (*α + β*)) and mean inputs are held fixed, Figure 5e. Thus, categorical distinctions drawn by statistical models that delineate and interpret step- versus ramp-like response curves as supporting binary or graded evidence accumulation^32^ may not be meaningful in dynamical neural models of noisy choice behavior, at least on the level of individual neural responses during the decision period ^66^.

In experiments, the decision threshold or level of activity right at the decision time of the winning choice neurons increases under pressure to respond rapidly^24^. This result is counterintuitive from the perspective of AB models, which increase speed by lowering the bound^24^. In WTA networks, the asymptotic activity of the winning neuron is proportional to 1/(1 – *α*), while speed increases with *α + β*. Starting from parameters consistent with a high reward rate, a speed-up can be achieved by increasing *α* or *β* or both (Figure S6a). Thus, except in the special case where only inhibition is allowed to increase, the activity of the winning neurons is predicted to increase under speed pressure, Figure 5f, as observed^24^.

By contrast, the decision threshold or final activity level of the winning choice neurons remains fixed in the strong-inhibition WTA models as *N* increases if parameters are held fixed, Figure 5g, as observed in experiment^9^.

Finally, in addition to self-adjustment as *N* changes, the nWTA network self-adjusts when the signal-to-noise ratio of the input decreases, taking longer to converge (Figure 4h) and averaging its inputs for longer during this extended convergence time (Figure S5g) both when the gap between the correct and incorrect options shrinks or the input noise increases. This self-adjustment capability is the result of recurrent inhibition, which may offer powerful computational flexibility to a decision-making circuit.

It has recently been shown in experiment that decision circuits can be trained to adapt their integration time to the time-varying statistics of the input^48^. The present result shows how such neural circuits, if similar to the nWTA network, may be able to automatically and instantly, without plasticity, adjust to the input statistics.

## Discussion

We have examined the efficacy of parallel computation for finding the best option when inputs are noisy, both in the abstract and in neural circuit models with leaky neurons. **max, argmax** are elemental operations in inference, optimization, decision making, action selection, consensus, error-correction, and foraging computations; we show that the nWTA neural network can accurately determine and report **max, argmax** in the noiseless and noisy input cases, with a computation time that meets the parallelism benchmarks (constant decision time in the noiseless case and a time that grows as *O*(log(*N*)) in the presence of noise).

When applied to psychophysical decision-making tasks^9;25;44;56^ (*N* ≤ 10), the model provides a explanation for Hick’s law^25^. We believe this is the first demonstration of Hick’s law within a neural network decision making model with self-terminating dynamics. Moreover, the model reproduces Hick’s law without the need to tune or change any parameters as the number of options changes or the task is made more or less difficult by varying the gap between options or the noise levels.

The flexible, self-adjusting nature of the neural WTA computation is due to recurrent inhibition, which automatically slows network convergence as the number of options are increased, noise is increased, or the gap between options is decreased; the additional threshold on inhibition in the nWTA circuit ensures that the WTA state exists in the presence of noise even in large networks.

Our work additionally reproduces a number of (sometimes counterintuitive) psychophysical and neural observations, including faster performance on correct than error trials, a higher pre-decision neural activity level when subjects are pressured to make faster decisions, and a natural tendency to weight early information over late (however, the extent of this tendency to impulsivity is tunable).

The efficiency of accurate parallel computation by the nWTA network holds not only in the regime of psychophysical decision making, but also when the number of options ranges in the thousands, corresponding possibly to a microscopic circuit inference operation of finding the maximally active neuron in a circuit. In this way, our work provides a single umbrella under which systems neuroscience questions about parallel computation distributed across large numbers of individual neurons and psychophysics questions about explicit decision making can be answered.

**Relationship to past work max, argmax** have been variously studied in computer science^3;14^ and in artificial^62^ and biologically plausible neural networks^18;22;23;28;35;40;55;62;67;69;70^. In some models, a nonlinear inhibitory contribution in the quite different form of a collective, network-level multiplicative (shunting) inhibition^70^ can produce fast convergence in the noise-free setting for small-to-medium *N*, but results in high sensitivity to fluctuations and thus a failure of accurate WTA across all values of *N*.

The leaky competing accumulator model^43;64^ is mathematically equivalent to the conventional WTA model, if *τ_η_* = *τ*^45^ and if the asymptotic state of the dynam-ics is used as decision criterion. Instead, however, the works use the crossing of a pre-determined threshold not tied explicitly to the asymptotic states as the decision criterion. In general, the works above do not consider WTA breakdown in the presence of noise and how decision time will scale at fixed accuracy in the noisy setting, or compare circuit dynamics with that seen in neurons during decision making. Neural network decision-making models make correspondences with neural phenomena^19;52;68^, but do not tend to systematically consider dynamics for multiple options.

Our results build on this existing work, extending it along several directions while unifying many previous results in the context of a single, neurally plausible network model.

### Biological mechanisms for thresholding inhibitory contributions

In circuits with separate excitatory and inhibitory neurons^12^, there are multiple candidate mechanisms for the inhibitory nonlinearity required by nWTA. These can be categorized by whether inhibitory interneurons are selectively tuned to particular principal cell inputs, or non-selective because they pool inputs from many principal cells. If inhibitory neurons are selective, then a simple threshold nonlinearity in the input-output transfer function, like the type-II firing rate responses in inhibitory neurons^26^, is sufficient. A similar effect could be achieved by fast-spiking inhibitory interneurons that act as coincidence detectors rather than integrators ^2^: these cells would respond weakly to low firing-rate inputs and reliably for high-rate inputs, thus effectively thresholding activity. Finally, if interneurons target pyramidal cells dendrites, then dendritic nonlinearities^37^ could threshold inhibitory input.

If inhibitory neurons are non-selective, then the nonlinearity must be present in the excitatory-to-inhibitory synapse so that the drive from the low-activity input principal cells is specifically ignored. If the excitatory to inhibitory synapses have low release probability and are strongly facilitating, only high firing rate inputs would make it through^71^.

Finally, while we have considered one particular form of the additional nonlinearity (i.e., an activation threshold), it may be possible to replace the activation threshold with other nonlinearities in either the excitatory or inhibitory units.

### Spatial organization of WTA circuit

Neural WTA models can be viewed as *N* principal cells or groups inhibiting each other (e.g. via interneurons private to each cell or group), which requires ~ *N*^2^ synapses. This connectivity is both dense and global (Figure 1c). Alternatively, all cells can drive a common inhibitory neuron (pool) which inhibits them all, an architecture that requires only ~ *N* synapses, a much sparser connectivity (only *O*(1) synapse per neuron) that is still global (Figure 1c). It may be possible to replace the global inhibitory neuron by local inhibitory neurons that pool smaller excitatory groups. However, local inhibition generically produces pattern formation^13^ and consensus formation with local connections can be unstable^4^, thus it is an interesting open question whether WTA can be implemented with purely local connectivity.

### Extensions and generalizations

Distributed decision making is a feature of many collective systems, including bacterial quorum sensing^46^, foraging and house-hunting in ants and bees ^5;16^, social and political consensus formation^36^, and economic choice behaviors. While our model is based on neural dynamics, the ingredients (self-amplification; recurrent nonlinear inhibition) are simple and should have analogues in other distributed decision-making systems. Our results suggest a scaling of *O*(log(*N*)) with the number of options and the existence of a thresholded or otherwise nonlinear inhibition if *N* is large.

Real world systems are bandwidth limited: neurons communicate with spikes; scout insects achieve consensus through brief interactions with subsets of others^15;16^. Here we have assumed high bandwidth communication: neurons exchanging analog signals in continuous time. Nevertheless, the principal cells in our model do not communicate their individual activation levels to all other cells; other principal cells receive information only about global activity in the network in the form of a single inhibitory signal, and there is noise, both forms of limited communication. In this sense, our results should generalize to the lower-bandwidth case. Studying the impact of low-bandwidth communication on WTA and parallel decision making in more detail is an interesting direction for future work.

## Methods

### Network model and dynamics

We consider *N* coupled neurons with activations *X_i_, i* ∈ {1,…, *N*}, and dynamics given by

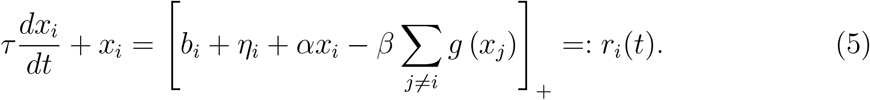

The neural nonlinearity is set to be the threshold-linear function: [*x*]_+_ = *max*[0,*x*]; *τ* is the neural time constant, *α* is the strength of self-excitation and *β* is the strength of global inhibition. *g(x)* is the inhibitory activation function: if *g* ≡ 1 the activation is fully linear and conventional WTA dynamics as studied in^69^ is recovered. We also consider an alternative, threshold-linear activation function *g*(*x*) = [*x* – *θ*]_+_ with threshold *θ*. We call the respective dynamical system *nWTA-dynamics*. The RHS of Equation (5) may be viewed as the instantaneous firing rate *r_i_(t)* of neuron *i*.

Each neuron receives a constant external drive *b_i_*. Neurons are ordered such that *b*_1_ > *b*_2_ > … > *b_N_*, and it is assumed that each input drives exactly one neuron, see Figure 1c. In addition, each neuron receives a private zero-mean fluctuation term *η_i_(t)*, which is modeled by statistically identical Ornstein-Uhlenbeck processes, i.e.,

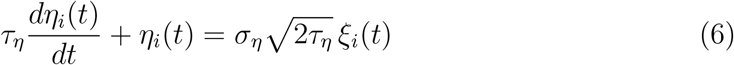

with Gaussian white noise *ξ_i_*(*t*), such that, ⟨*ξ_i_*(*t*)⟩ = 0, ⟨*ξ_i_*(*t*)*ξ_j_*(*t*’)⟩ = *δ_ij_·δ*(*t* – *t*’). It follows that ⟨*η_i_(t)*⟩ = 0 and 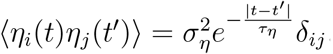.

A network that converges to a unique WTA state with non-noisy inputs and internal dynamics need not do the same when driven by noise. Noise kicks the state around and the system generally cannot remain at a single point. Nevertheless, the network state can still flow toward and remain near a fixed point of the corresponding deterministic system (S2.5, Figure S3a-d). We will refer to such behavior in the noise-driven WTA networks as successful WTA dynamics, defined in terms of one neuron reaching a criterion distance from the deterministic WTA high-activity attractor (set by the dynamical system, not an external threshold) while the rest are strongly suppressed.

### Conditions for WTA dynamics

Analysis of the linear stability of the noise-free conventional dynamical system Eqn.(5) with *g* ≡ 1 (see S2.1 and^69^) reveals one eigenvalue *λ*_w,hom_ = *α* – (*N* – 1)*β* with uniform eigenvector 1 = (1,…, 1)^τ^, and an (*N* – 1)-fold degenerate eigenvalue *λ*_W,diff_ = (*α* + *β*) whose eigenvectors are difference modes with entries that sum to zero. If *α*+*β* > 1 the difference modes grow through a linear instability, and the eventual (non-trivial) steady states involve only one active neuron. If *α* < 1 and *β* > 1 – *α*, the network will always select a unique winner for each b and initial condition^69^. For a discussion of more general constraints on *α,β*, see S2.1.

After meeting the conditions for stability and uniqueness (*α* < 1,*β* > (1 – *α*)), there is freedom in the choice of how the strength of global inhibition *β* scales with *N*: We may choose *β* ~ *O*(1), which we call the *strong inhibition* condition, or *β* ~ *β_0_/N*, the *weak inhibition* condition. In the weak inhibition regime, we set 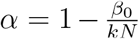 (with *k* > 1; specifically, we choose *β*_0_ = 1, *k* = 2 throughout the paper) for stability.

For simplicity, throughout the paper we consider the case where all neurons start at the same resting activity level x(0) = (*x*_0_,…, *x*_0_)^τ^. In this case, the winner is the neuron with the largest input. (For heterogeneous initial conditions the situation is more complex, since the wrong neuron can be pushed to be the winner just by starting at large enough activity to suppress all other neurons, see also the discussion in^69^.) We further assume *x*_0_ = 0 (if *x*_0_ > 0 there is an initial transient that scales logarithmically with *N* but is unrelated to the actual WTA-computation, see S2.2).

In the case of noisy conventional WTA dynamics (but not for the nonlinear WTA model) the asymptotic activation of the winner 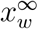 depends on the number of neurons and the noise amplitude in a nontrivial manner (see S2.5). For convenience we thus define *T*_WTA_ as the time some neuron reaches an activation level greater than or equal to 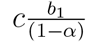 with c ≲ 1 (we use *c* = 0.8) in all simulations (noisy or deterministic). We emphasize that this convergence criterion is nonetheless set by the dynamics and hence inherently different from an external arbitrary threshold, and does not change the scaling of either the deterministic or noisy dynamics, see S2.2.

### Simulations and analysis

All simulations and analyses were carried out using standard Python packages (Python 2.7.12, NumPy 1.11.0, SciPy 0.17.0). The dynamics Eqn. (5) was solved by simple forward-Euler integration with integration time step Δ*t* ∈ [10^-5^*τ*, 0.2*τ*] depending on numerical requirements. For the simulation of Ornstein-Uhlenbeck noise we made use of exact integration on a time grid with increment Δ*τ*^20^, i.e.,

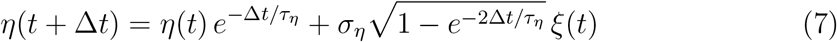

