## Supplementary Material for "Robust parallel decision-making in neural circuits with nonlinear inhibition"

### Supplementary Information

**S2 Winner-take-all networks 5**
S2.1 Allowed $\alpha$ values for strong and weak inhibition in determini-stic WTA networks 5
S2.2 Derivation of $T_{\mathrm{WTA}}$ for deterministic rate dynamics and $\mathbf{b} = (b + \Delta, b, \ldots, b)^\top$ 6
S3.1.2 WTA-performance and Hicks law for fixed network parameters $\alpha, \beta$ . 18

### S1 Scaling of noisy argmax: sequential algorithm and parallelism benchmark

#### S1.1 Scaling of time for fixed accuracy

We first compute how long to observe each input in the sequential comparison process to obtain a desired error.

**N log(N) decision time is sufficient to maintain high fixed accuracy** Assume there are $N$ noisy inputs with means $b_1 > b_2 \ldots \geq b_N$ and some bounded variances. Observing the inputs for unit time yields random variables $\tilde{b}_1, \ldots, \tilde{b}_N$, also with means $b_1, \ldots, b_N$, and bounded variance. Next assume that we observe these variables for a number of time steps $T$ (defined in terms of the unit time). It will be convenient if samples at different time steps are independent or weakly correlated, and for reasonable noise models this can be arranged by making the unit time sufficiently large.

Consider the differences $\delta_i = \tilde{b}_1 - \tilde{b}_i$, $(i = 2, \ldots, N)$, which have means $\Delta_i = b_1 - b_i > 0$ and some variances $\sigma_i^2$. We consider the time-averages of these observations,

$$\bar{\delta}_i^T = \frac{1}{T} \sum_{t=1}^{T} \delta_i^{(t)},$$

where $\delta_i^t$ is the $t^{th}$ observation of the $i^{th}$ difference. (It is equivalent to first average the $\tilde{b}_i$'s over $T$ then take the differences). We may write $\bar{\delta}_i^T = \delta_i + \xi_i^T$, where $\mathbb{E}[\bar{\delta}_i^T] = \Delta_i$ and $\xi_i^T$ is a zero-mean random variable such that $\mathrm{Var}[\bar{\delta}_i^T] = \langle (\xi_i^T)^2 \rangle = \sigma_i^2/T$.

The wrong input is identified by **argmax** if any $\bar{\delta}_i < 0$. Therefore,

$$P(\text{error}) \leq \sum_{i=2}^{N} P(\bar{\delta}_i^T < 0). \tag{1}$$

As $T$ increases, $\bar{\delta}_i^T$ concentrates around its mean $\Delta_i > 0$, with typical fluctuations shrinking as $1/\sqrt{T}$. We assume that $T$ is sufficiently long that the probability of any particular error is small; this will be true for any reasonable total error probability. Thus the event $P(\bar{\delta}_i^T < 0)$ lives in the tail of this distribution, outside the range of typical fluctuations. For a large class of distributions, the probability of these large deviations falls off exponentially with $-\Delta_i^2 T$ in the exponent (see end of this section).

The individual error probabilities in Equation (1) decay exponentially strongly with $T$ and $\Delta_i$. The largest term involves the topmost gap, $\Delta_1$. If this gap is held fixed as $N$ is scaled up, the remaining error terms are each smaller but there are $N$ such terms, producing a total error of the form $e^{-\Delta_1^2 T}(1 + (N-1)\epsilon)$, where $\epsilon < 1$. To obtain a total error that does not scale with $N$, it follows that the additional time-complexity of the computation per input is bounded by $T \sim (\log(N)/\Delta_1^2)$ for noisy inputs. Defining $\Delta_1 \equiv \Delta$ for notational convenience, the time-complexity of noisy **max** for fixed $\Delta$ is $T_{\mathrm{S}} = NT \sim O(N \log(N)/\Delta^2)$.

If all the inputs $b_i$ are drawn uniformly from a random interval, the gap $\Delta$ itself scales as $1/N$. Then the time-complexity of noisy **max**, **argmax** with fixed error probability scales as $T_{\mathrm{S}} \sim N^3 \log(N)$, in contrast to the noise-free scaling in this case of $T_{\mathrm{S}} \sim N$.

**N log(N) decision time is necessary to maintain high fixed accuracy for Gaussian inputs** Equation 1 is an upper bound on the decision time, both because of the use of a union bound to bound the error probability and because the exponential tail bound on fluctuations may not be tight, though it often is (exponential tail bounds are described further at the end of this section).

A more accurate but less intuitive result can be obtained by examining the distribution of the maximum of the set of random variables $\{\bar{\delta}_i^T\}$, which can usually be described by an extreme value distribution[2]. For Gaussian random variables, the maximum concentrates around the mean of this distribution, which scales as $\sigma\sqrt{\log(N)}$[2]. In order to maintain a fixed probability that $\tilde{b}_1$ is greater than this value, we must remove the $N$-dependence, and consequently $\sigma$ must scale as $1/\sqrt{\log(N)}$. Equivalently, the observation time $T$ should scale as $\log(N)$ (recall that $\sigma$ goes as $1/\sqrt{T}$).

While the above result is described for Gaussian random variables, the variables $\{\tilde{b}_2, \ldots, \tilde{b}_N\}$ are the sums of independent random samples and are thus Gaussian under mild conditions (by the Central Limit Theorem). Thus the scaling time will be $N \log(N)$ in many cases.

We also confirm these theoretical arguments numerically, in Figure 1b. For these plots we integrate $N$ independent inputs generated by Ornstein-Uhlenbeck processes with the first input having a mean of 1 and the remainder having means of 0.95, and all inputs having standard deviation of 0.2. Results are equivalent for independent discrete samples from Gaussian distributions (the choice of O-U processes is simply to make the continuous time process mathematically well-defined).

#### S1.2 Error probability at fixed observation time

We next fix the observation time $T$, and examine the maximum error probability. For the quasi-2D case, the largest input has mean $b_1 = b + \Delta$, and the remaining $N - 1$ inputs have mean $b$. For convenience we set $b = 0$ (as this simply shifts the origin).

We consider the case where noise is Gaussian, so that the random variables $\tilde{b}_i$ are drawn from a normal distribution with means $b_i$ and variance $\sigma^2/T$. As in the previous section, we consider the distribution of the maximum of the set of random variables $\{\tilde{b}_2, \ldots, \tilde{b}_N\}$, and note that this concentrates around its mean, which scales as $\sigma\sqrt{\log(N)}$[2]. Call this maximum $\mu_{max}$.

The probability of error is then $P(\tilde{b}_1 < \mu_{max}) = \Phi((\mu_{max} - \Delta)/\sigma) = \Phi(k_1\sqrt{\log(N)} + k_2)$, where $k_1$ and $k_2$ are constants that depend on $\Delta$ and $\sigma$, and $\Phi$ is the cumulative distribution function of a Gaussian. Equivalently, $p_{error}$ is a sigmoidal function of $\sqrt{\log(N)}$. In the tail of the distribution $1 - \Phi$ can be approximated by $e^{-x^2}/x$[4], yielding an approximately inverse power law dependence of error probability on $N$.

The decay of accuracy with $N$ for fixed time in the WTA network is also well-fitted by $\Phi(k_1\sqrt{\log(N)} + k_2)$ except at small $N$.

#### S1.3 Exponential decay of $P(\bar{\delta}_i^T < 0)$

We show here that the probability $P(\bar{\delta}_i^T < 0)$ falls off exponentially in $T$ for two cases that cover a very wide set of possible distributions on the inputs $b_i$.

First, if the $\bar{\delta}_i^T$'s are bounded, as is common for biological variables, then the result follows from Hoeffding's inequality [5].

Second, note that the $\bar{\delta}_i^T$'s are sums of random observations and thus the Central Limit Theorem should apply. If the $\bar{\delta}_i^T$'s are Gaussian, then the exponential fall-off of $P(\bar{\delta}_i^T < 0)$ follows from the asymptotic form of the complementary error function [1] (see below). The results are likely to apply more generally, for example through the Chernoff bounds on the tails of other distributions.

**Bounded variables.** Assume that each $\bar{\delta}_i^T$ comes from a distribution bounded in $[a, b]$, with $b - a = R$, and mean $\Delta_i$.

$$\begin{aligned} P(\bar{\delta}_i^T < 0) &= P(\bar{\delta}_i^T - \Delta_i < -\Delta_i) \\ &\leq \exp\left(-\frac{2\Delta_i^2 T}{R^2}\right). \end{aligned} \quad (2)$$

Here the inequality comes from applying Hoeffding's inequality [5].

Now let $T_i = k_i \log(N)$, with $k_i \geq R^2/(2\Delta_i^2)$, and choose the observation time $T$ to be $\max_i T_i$.

**Normal distribution.** Assume that each $\bar{\delta}_i^T$ is normally-distributed, so that $\bar{\delta}_i^T \sim \mathcal{N}(\Delta_i, \frac{\sigma_i^2}{T})$. Consider the standardized variable

$$W_i = \frac{\bar{\delta}_i^T - \Delta_i}{\sigma_i/\sqrt{T}},$$

which has zero mean and unit variance.

$$\begin{aligned} P(\bar{\delta}_i^T < 0) &= P\left(W_i < -\frac{\sqrt{T}\Delta_i}{\sigma_i}\right) \\ &\leq \exp\left(-\Delta_i^2 T/(2\sigma_i^2)\right), \end{aligned} \tag{3}$$

where the inequality follows from a loose bound on Gaussian tails [4].

Now let $T_i = k_i \log(N)$, with $k \geq (2\sigma_i^2)/\Delta_i^2$ and choose $T$ to be $\max_i T_i$.

#### S1.4 More complex/adaptive strategies

We have derived the parallelism benchmark above by considering independent integration of inputs in parallel. A natural question is whether a more complex adaptive strategy could outperform this independent parallel integration, taking less time to achieve a desired accuracy level by comparing the values across inputs and terminating the computation using a more complex criterion rather than waiting a fixed time or integrating to a fixed threshold. We do not consider such strategies in the main text, for two reasons.

First, more complex algorithms require comparing the alternatives at each step. To compare even a fraction of the $N$ alternatives requires an additional time complexity of $\sim N$ at each step. Thus while these algorithms may use fewer samples of input, the total time complexity will be greater than the $O(\log(N))$ parallelism benchmark derived above.

Second, we consider below a particular example of a more complex strategy that is known to be optimal in the limit of vanishing error rate. We show that even if we neglect the additional complexity of comparing the inputs at each step, the integration time for this more complex strategy still grows as $\sim \log(N)$ and there is no advantage over the parallelism benchmark we have derived above. Thus, this benchmark is likely to be optimal.

**The diff-AB strategy** There is no simple decision model (i.e., there is no simple approximation to the full Bayesian expression) known to be optimal for multi-AFC tasks with more than two alternatives [3, 6, 7]. One candidate algorithm considers the difference between the largest and second-largest integrated alternative, and terminates the decision when this difference crosses a threshold [6]. This strategy (henceforth diff-AB) requires identification of the top two integrated alternatives at each time, which requires traversing the list of options, adding an additional time-complexity of $N$ beyond that of the simple integration in parallel. However, if the additional time complexity of the comparison is neglected and only the integration time of each option is considered, then diff-AB is optimal in the limit of vanishing error rate[6].

We find numerically that the decision time required for diff-AB to reach a decision with fixed accuracy grows as $\sim \log(N)$ (ignoring the time to compare options at each step), Figure S2a. Thus, the scaling of diff-AB is not an improvement over the parallelism benchmark.

In Figure 5, we show that for small values of $N$, a self-terminating WTA network is able to quantitatively outperform simple parallel integration (note that the scaling remains $\sim \log(N)$, so there is no contradiction or violation of a theoretical bound). In Figure S6b we show that WTA is competitive with the integration time required by diff-AB for low error (again, ignoring the time to compare options at each step).

Since diff-AB is known to be optimal for vanishing error[6], WTA networks not only show optimal scaling but can approach quantitatively optimal decision times.

### S2 Winner-take-all networks

#### S2.1 Allowed $\alpha$ values for strong and weak inhibition in deterministic WTA networks

We consider networks described by the equation (Eqn.(1) in the main text):

$$\tau \frac{dx_i}{dt} + x_i = \left[ b_i + \eta_i(t) + \alpha x_i - \beta \sum_{j \neq i} x_j \right]_+ \tag{4}$$

The coupling matrix $W$ has one eigenvalue $\lambda_{\mathrm{W,com}} = \alpha - (N-1)\beta$ with uniform eigenvector $\mathbf{1} = (1, \ldots, 1)^\top$, and an $(N-1)$-fold degenerate eigenvalue $\lambda_{\mathrm{W,diff}} = (\alpha + \beta)$ whose eigenvectors are difference modes with entries that sum to zero. Assuming that the neurons do not encounter the rectification threshold, the network equation is linear and has one eigenvalue at $\lambda_{\mathrm{com}} = -1 + \alpha - (N-1)\beta$, and $N-1$ eigenvalues of size $\lambda_{\mathrm{diff}} = -1 + \alpha + \beta$. WTA dynamics (i.e. competition) requires that the difference modes are unstable [8], so that

$$-1 + \alpha + \beta > 0. \tag{5}$$

For the activity levels to remain bounded (stability), either $\alpha < 1$ or a saturating nonlinearity should be included in the neural transfer function. If $\alpha < 1$, then the network dynamics remain linear until they encounter the rectification threshold where the combination of unstable dynamics and the threshold keeps the network at a steady-state. This can be visualized on an energy landscape, with the network state moving to lower energies (shown for $N = 2$ in Figure S1a,b).

Below we analyze the existence of the WTA state for different scalings of $\beta$ with N.

**Weak inhibition ($\beta = \beta_0/N$)** WTA dynamics in a weakly inhibiting network requires that $\alpha \to 1$ as $N \to \infty$. To see this, first consider the case $\alpha < 1$. According to Equation 5, $\alpha > 1 - \beta_0/N$. Thus, we have $1 - \beta_0/N < \alpha < 1$, and as $N \to \infty$ it follows that $\alpha \to 1$. Next, consider the case $\alpha > 1$ (with some saturation assumed so that the system is stable). Suppose all but two neurons are driven down to some baseline state $x_{base}$, which can be 0.

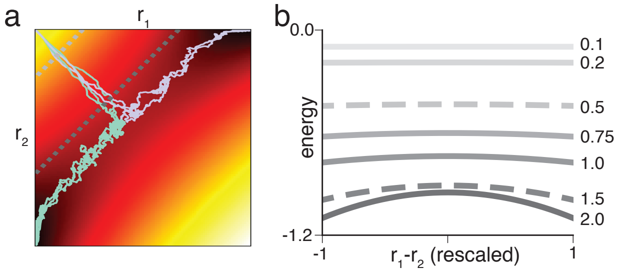

Figure S1: **Fixed-point dynamics of WTA** (a) Sample trajectories from different trials of a network with $N = 2$ neurons, $\alpha = 0.55$, $\beta = 0.6$. Firing rates of neurons shown on $x$ and $y$ axes. Background colors indicate the energy of the corresponding state (dark = low energy). (b) Slices of the energy landscape shown in (a). Slices taken along lines of the form $r_1 + r_2 = k$ (dashed lines correspond to dashed lines in (a)). Numbers on the right indicate the corresponding $k$. Note that the energy landscape is near-flat for small values of $k$, allowing good integration.

The dynamics can be written as:

$$\begin{aligned}\dot{x}_1 &= (\alpha - 1)x_1 - \frac{\beta_0}{N}x_2 - \frac{N-2}{N}\beta_0 x_{base} + b_1 + \xi_1 \\ &\approx (\alpha - 1)x_1 - \beta_0 x_{base} + b_1 + \eta_1,\end{aligned}$$

with a similar equation for $x_2$ (the approximation is for large $N$). In this case $x_1$ and $x_2$ are effectively uncoupled and since $\alpha > 1$, both can both become "winners" for sufficiently large $b_1, b_2$ (that do not scale up with $N$). Thus, for a unique winner, the strength of self-excitation must remain $\alpha < 1$.

In simulations, to satisfy $1 - \beta_0/N < \alpha < 1$, we set $\alpha = 1 - \beta_0/kN$, where $k > 1$ is some constant.

**Strong inhibition ($\beta = \beta_0$, independent of $N$)** If the losing neurons are 0, as in the deterministic network, then the WTA state exists. However, if $x_{base} > 0$ or shrinks to zero more slowly than $1/N$, this network does not have a winner. Suppose $N - 1$ neurons are at $x_{base}$ and the remaining neuron (say neuron 1) is active. Then

$$\dot{x}_1 = -(1 - \alpha)x_1 - \beta_0(N - 1)x_{base} + b_1 + \eta_1(t), \tag{6}$$

with steady-state $x_1 = \frac{-\beta_0(N-1)x_{base}+b_1}{1-\alpha}$. For large $N$, if $\alpha < 1$ this steady-state is stable and negative, so $x_1$ is attracted to 0. If $\alpha > 1$ the steady-state is unstable, positive and large, so $x_1$ is driven away from it and towards 0. In both cases the network does not show a winner.

#### S2.2 Derivation of $T_{\mathbf{WTA}}$ for deterministic rate dynamics and $\mathbf{b} = (b + \Delta, b, \ldots, b)^\top$

The solution of the deterministic rate dynamics Equation(4), $\eta_i(t) \equiv 0$ for all $i$, with homogeneous initial conditions $\mathbf{x}_0 = (x_0, \ldots, x_0)^\top$ and "two-dimensional" drive $\mathbf{b} = (b+\Delta, b, \ldots, b)^\top$

can be derived in a piecewise manner by taking into account at which times the threshold linearity returns either zero or positive values. For the sake of simplicity, we assume $\mathbf{x}_0 \equiv \mathbf{0}$, i.e., the neurons start the WTA competition from a state of zero activation[1]. The dynamics is then given by

$$\frac{dx_w(t)}{dt} + x_w(t) = b + \Delta + \alpha x_w(t) - \beta(N-1)x_\ell(t) =: r_w(t) > 0 \tag{7a}$$

$$\frac{dx_\ell(t)}{dt} + x_\ell(t) = b + \alpha x_\ell(t) - \beta(N-2) - \beta x_w(t) =: r_\ell(t) > 0 \tag{7b}$$

The solutions $x_w(t), x_\ell(t)$ are

$$\begin{aligned} x_\ell(t) = \frac{1}{N\lambda_{\text{diff}}\lambda_{\text{com}}} \times \Big( & -Nb\lambda_{\text{diff}}\left(1 - e^{-t(-\lambda_{\text{com}})}\right) \\ & - \Delta\big(N\beta - \lambda_{\text{diff}} e^{-t(-\lambda_{\text{com}})} + \lambda_{\text{com}} e^{-t(-\lambda_{\text{diff}})}\big)\Big) \end{aligned} \tag{8}$$

$$\begin{aligned} x_w(t) = \frac{1}{N\lambda_{\text{diff}}\lambda_{\text{com}}} \times \Big( & - N\big(b\lambda_{\text{diff}} + \Delta(\lambda_{\text{com}} - \beta)\big) \\ & + (Nb + \Delta)\lambda_{\text{diff}}\, e^{-t(-\lambda_{\text{diff}})} + (N-1)\,\Delta\,\lambda_{\text{diff}}\, e^{-t(-\lambda_{\text{diff}})}\Big) \end{aligned}$$

where $-\lambda_{\text{diff}} = 1 - (\alpha + \beta)$.

We define $T_{\text{WTA}}$ as the time $t_0$ when the rate $r_\ell(t)$ of the $N-1$ loser neurons crosses zero from above, i.e., $b + \alpha x_\ell(t_0) - \beta(N-2) - \beta x_w(t_0) \overset{\downarrow}{=} 0$. This $t_0$ can be computed numerically.

**Scaling for $\beta, \alpha \sim$ const** Inserting Eqns. (8), and assuming $N \gg 1$, we obtain the condition for the rectification to take action as

$$\begin{aligned} & b(-\lambda_{\text{diff}}) - \beta\Delta\left(-\lambda_{\text{diff}} e^{-t(-\lambda_{\text{diff}})}\right) \overset{\downarrow}{=} 0 \\ \Leftrightarrow \quad & T_{\text{WTA}} \sim \frac{\log\left[\frac{\beta\Delta - b(-\lambda_{\text{diff}})}{\beta\Delta(\alpha+\beta)}\right]}{\lambda_{\text{diff}}} \sim \log\left[1 - \text{const} \times \frac{b}{\Delta}\right] \end{aligned} \tag{9}$$

The time $T_{w,\epsilon}$ it takes the winner to reach any $\epsilon$-distance to its asymptotic state $x_w^\infty = (b + \Delta)/(1 - \alpha)$, starting from some value $x_w(T_{\text{WTA}})$ the moment the last competitor is rectified, is independent of $N$, so does not add any further $N$-dependence:

$$T_{w,\epsilon} = -\frac{1}{1-\alpha} \log\left[\frac{(b+\Delta)(1-\epsilon)}{(b+\Delta) - x_w(T_{\text{WTA}})(1-\alpha)}\right] \tag{10}$$

[1] If $0 < \frac{b}{(N-1)\beta - \alpha} < x_0$ there will be an initial transient phase where none of the neurons has suprathreshold input, and hence all $x(t)$ will have an exponential decay $x_{w/\ell}(t) = x_0\, e^{-t/\tau}$. The neuron with the strongest drive, i.e., $x_w(t)$, will become active first when its RHS crosses zero from below, i.e., at time $t_{w,0} = \log\left[\frac{x_0((N-1)\beta-\alpha)}{b}\right]$. Note that this logarithmic $N$-dependence is trivial and not related to the actual WTA-dynamics.

**Scaling for** $\beta \sim 1/N, \alpha \sim 1 - \frac{1}{2N}$ In this case we need to take into account that the coupling terms $\alpha, \beta$ carry an $N$-dependence themselves. Analysis of the input of $x_\ell$ reveals

$$\frac{2N^2(2\Delta + b) - (\Delta + bN)\, e^{-t(1-\frac{1}{2N})} - (4N^2 - 1)\Delta\, e^{t/2N}}{N(2N-1)} \overset{\downarrow}{=} 0 \tag{11}$$

$$\overset{N\gg 1}{\longrightarrow} \quad b - 2\Delta\left(e^{t/2N} - 1\right) \overset{\downarrow}{=} 0 \quad \Leftrightarrow \quad T_{\text{WTA}} \sim 2N \log\left[1 + \frac{b}{2\Delta}\right] \sim N$$

The time $T_{w,\epsilon}$, Equation (10), it takes the winner to reach any $\epsilon$-distance to its asymptotic state $x_w^\infty = (b+\Delta)/(1-\alpha) = 2N(b+\Delta)$ asymptotically scales $\sim N$, and thus again does not add further $N$-dependence:

$$T_{w,\epsilon} = -2N \log\left[\frac{(b+\Delta)(1-\epsilon)}{(b+\Delta) - \frac{x_w(T_{\text{WTA}})}{2N}}\right] \overset{N\gg 1}{\sim} 2N \tag{12}$$

To summarize, the decision time $T_{\text{WTA}}$ grows linearly with $N$ for deterministic (and noisy, see Figure S2c) input cases because the initial total inhibition at each neuron is $O(1)$, and is roughly canceled by the excitatory drive at each cell. The eventual winner and losers are thus somewhat isolated from each other, individually integrating their input drives with the slow effective time-constant $\sim \tau/(1-(\alpha+\beta))$. As $N$ increases, this time constant grows linearly with $N$. The individual integration process continues at length, until the losers and eventual winner finally separate enough that the nonlinear portion of WTA dynamics pushes them to their steady-state activations.

#### S2.3 Deterministic WTA-dynamics primarily determined by the top gap $\Delta$

In the main manuscript we focus on the case where all inputs $b_i$ are identical, apart from one that receives a positive bias $\Delta$. Here, we analyze how $T_{\text{WTA}}$ depends on the distribution of inputs.

First, in Figures S2b,c we show that in the absence of noise $T_{\text{WTA}}^{\text{uni}}$ (gray dashed), i.e., for fixed $\Delta = b_1 - b_2$ and uniform-randomly distributed inputs $b_{i\geq 3} \in \text{U}[0,1]$, is longer than $T_{\text{WTA}}^{\text{2d}}$ for quasi-2d inputs (black solid). This observation might suggest that the average activation of the $(N-2)$ bulk neurons should have some impact on $T_{\text{WTA}}$ for finite $N$. Figure S2c shows the results of simulations where the bulk $b_i, i \geq 3$ are drawn from uniform distributions $\text{U}[\mu - L/2, \mu + L/2]$ with varying mean $\mu \in \{0.1, 0.45, 0.8, 0.85\}$ (light to dark gray) and fixed width $L = 0.1$ for $\Delta = 0.1$.

For the interval with the largest mean this implies an upper bound $(1-\Delta)$. Surprisingly, only this interval has a measurable effect on the decision time, in that $T_{\text{WTA}}$ approaches the $N \to \infty$-limit of the effectively 2-dimensional case for large $N$ (Equation(9), dark gray line versus cyan line). Bulk inputs drawn from any of the other considered intervals have virtually no effect on $T_{\text{WTA}}$, and instead give rise to decision time scaling that equals that of the $N = 2$-case (red line).

An analogous result is obtained for a variation of bulk interval width $L \in \{0.18, 0.36, 0.9\}$ and fixed mean $\mu = \frac{1-\Delta}{2}$, Figure S2b. Again, only the last interval with upper bound given by $1-\Delta$ has any impact on $T_{\text{WTA}}$, in that it approaches Equation(9) for $N \to \infty$.

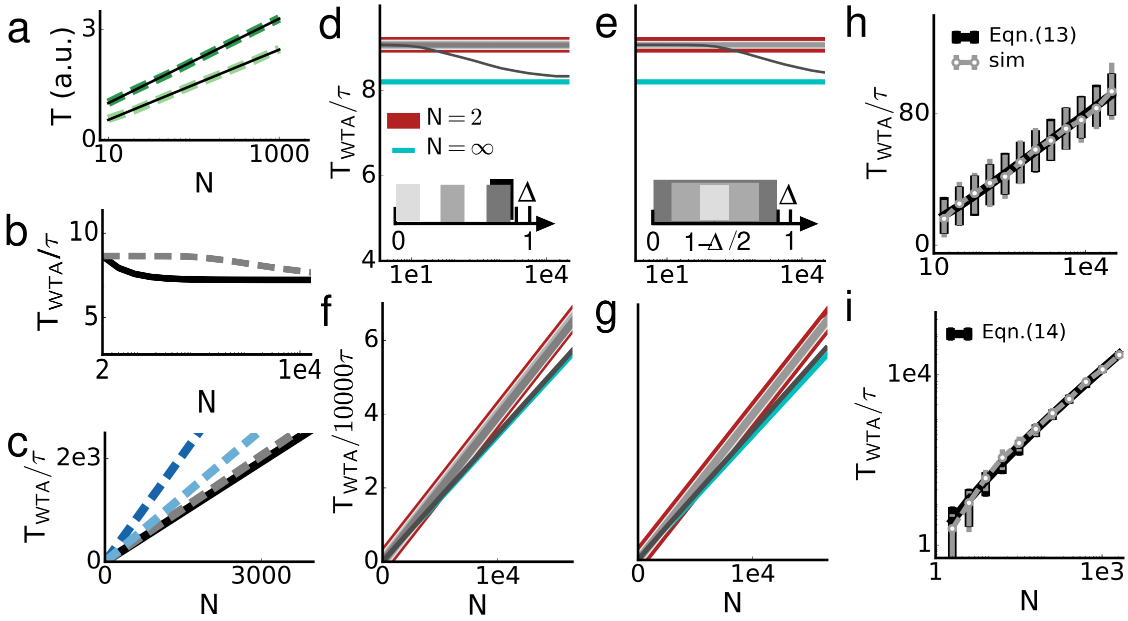

Figure S2: **Scaling of decision time and dependence on strategy and input drive distribution** (a) Scaling of a diff-AB strategy for two accuracy levels $A = 0.6$ (light green) and $A = 0.8$ (dark green) at noise amplitude $\sigma_\eta = 0.12$ and gap size $\Delta = 0.075$. (b) Scaling of $T_{\mathrm{WTA}}$ quasi-2d input (black) and inputs with fixed top-gap $\Delta$ and otherwise uniformly distributed inputs (dashed gray) for networks with strong inhibition. For quasi-2d inputs the results are analytically derived and exact. (c) Gray lines: same as (b) but for weak inhibition. Blue lines: scaling for same networks in the presence of noise ($\sigma_\eta = 0.6$). Dark blue: quasi-2d, light blue: fixed top gap and otherwise uniform inputs. (d–g) Scaling of deterministic $T_{\mathrm{WTA}}$ for fixed top gap $\Delta = b_1 - b_2$ and different distributions of $b_{i\geq 3}$; (d,e) strong inhibition, (f,g) weak inhibition. Gray lines correspond to distributions in inset; cyan: $N = \infty$ scaling; red: $N = 2$-scaling (note that for weak inhibition $\alpha$ still scales $\sim 1/N$, see text for details). (h) Deterministic $T_{\mathrm{WTA}}$-scaling is $\sim \log(N)$ for strong and (i) $\sim N\log(N)$ for weak inhibition if $\forall_i : b_i \in \mathrm{U}[0,1]$; black: theory, gray: simulations.

Identical results hold for weak inhibition $\beta \sim 1/N$, see Figure S2c,d, which leads to a $T_{\mathrm{WTA}}$-scaling linear in $N$, see Equation(11). The red line in Figure S2e,f is the solution for only two competing neurons ($N = 2$), but with inhibition that still scales as $\alpha = 1 - \frac{1}{2N}$, $\beta = \frac{1}{N}$, i.e., $T_{\mathrm{WTA}} = 2N \log\left[\frac{2(1-\Delta)}{3\Delta}\right]$.

Moreover, we note that these results are not limited to the choice of bulk intervals here, but are the same for other distributions for the bulk $b_i$, e.g., Gaussian or exponential: Only if there is a finite density close to the second largest input will there be a measurable impact on $T_{\mathrm{WTA}}$. In the limit case of quasi-2d the $N - 1$ competitors of the biased neuron have the highest activation, and hence strongest inhibitory effect on all others of all considered scenarios. That is why for deterministic dynamics the quasi-2d dynamics is the fastest, because suppression of losers is most effective. This is different for noisy inputs, see, e.g., blue curves in Figure S2c. The reason for this will be discussed in S2.5.

Together, these numerical findings suggest that for deterministic dynamics the decision time $T_{\text{WTA}}$ is primarily determined by the gap $\Delta$ between the two largest inputs, with an at most minor secondary contribution from—and little sensitivity to—the distribution of the rest of the inputs.

#### S2.4 Scaling of deterministic $T_{\text{WTA}}$ for uniformly distributed $b_i$

Now consider the case where all inputs are selected uniformly and randomly over the interval $[0, 1]$, such that the gap $\Delta$ is not held fixed as $N$ is varied. The observation that only $\Delta$ matters for $T_{\text{WTA}}$ allows to analytically compute the decision time, which now acquires a dependence on $N$: $T_{\text{WTA}}$ grows as $\log(N)$. This dependence on $N$ arises entirely from the shrinking size of the gap $\Delta$ between the top two inputs: The gap between $N$ uniformly randomly generated entries shrinks as $1/N$. Assume that for all $i \in \{1, \dots, N\}, b_i \sim \text{U}[0, 1]$ and without loss of generality, $b_1 > b_2 \geq \dots \geq b_N$. The resulting gap $\Delta := b_1 - b_2$ between the largest and second largest drive is then Beta-distributed, such that $\Delta \sim \text{Beta}[1, N]$. For large $N$ this distribution approaches the exponential distribution $P(\Delta) = N\, e^{-N\Delta}$. Motivated by the simulation results discussed above that it is predominantly the top gap that settles $T_{\text{WTA}}$, we can estimate mean and variance of $T_{\text{WTA}}^{\text{uni, strong}}$ by computing the respective averages over $\Delta$ for $T_{\text{WTA}}^{2d,\text{strong}}(\Delta)$:

$$\begin{aligned} \text{E}[T_{\text{WTA}}^{\text{uni,strong}}] &= \int_0^\infty T_{\text{WTA}}^{2d,\text{strong}}(\Delta) N e^{-N\Delta}\, d\Delta \qquad (13) \\ &= \frac{\gamma + \log\left(\frac{N(\alpha+\beta-1)}{\beta(\alpha+\beta)}\right) - e^{-\frac{N(-\lambda_{\text{diff}})}{1-\alpha}} \text{Ei}\left[\frac{N(-\lambda_{\text{diff}})}{1-\alpha}\right]}{\alpha+\beta-1} \overset{N \gg 1}{\sim} \frac{\gamma + \log\left(\frac{N(\alpha+\beta-1)}{\beta(\alpha+\beta)}\right)}{\alpha+\beta-1}, \end{aligned}$$

where we for simplicity chose $b = 1 - \Delta$ to define a scale. $\gamma \approx 0.5772$ is Euler's constant, and $\text{Ei}[x] = -\int_{-x}^{\infty} \frac{e^{-t}}{t}\, dt$. Even though for valid choices of $\alpha$ and $\beta$ the term $e^{-\frac{N(-\lambda_{\text{diff}})}{1-\alpha}}$ grows exponentially in $N$, the Ei[.]-term decreases even faster, and their product can thus be neglected for large $N$, leaving only the logarithmic $N$-dependence of $T_{\text{WTA}}$. In the same way we can compute the variance $\text{Var}[T_{\text{WTA}}^{\text{uni,strong}}] = \int_0^\infty T_{\text{WTA}}^{2d,\text{strong}}(\Delta)^2 N e^{-N\Delta}\, d\Delta - \text{E}[T_{\text{WTA}}^{\text{uni,strong}}]^2$, and comparing both to our simulation results proves a striking agreement, see Figure S2g. Analogously, for weak inhibition we obtain (see Figure S2h)

$$\text{E}[T_{\text{WTA}}^{\text{uni,weak}}] \sim \int_0^\infty T_{\text{WTA}}^{2d,\text{weak}} N e^{-N\Delta}\, d\Delta \sim 2N\left(\gamma + \log\left(\frac{N}{2}\right)\right) \sim N \log(N)\,. \qquad (14)$$

Both for weak and strong inhibition the scaling of deterministic WTA for a shrinking gap is thus highly unfavorable compared to the serial performance ($\text{T}_p \sim$ const) which does not depend on the size of the gap in any way. However, as we will see, in the noisy case, the dependence of integration time on the gap will prove useful, allowing the network to self-adjust as the task becomes more difficult.

#### S2.5 Self-consistent solution for stationary states of noisy WTA networks

We assume $\tau_\eta \ll \tau$, such that the timescale of the noise fluctuations is much faster than the fluctuations around the steady state activities $\bar{x}_w, \bar{x}_\ell$. Considering again $\mathbf{b} = (b+\Delta, b, \ldots, b)^\top$ we define

$$\begin{aligned} \mu_w(\bar{x}_w, \bar{x}_\ell) &:= (\alpha\bar{x}_w - \beta(N-1)\bar{x}_\ell + b + \Delta) \\ \mu_\ell(\bar{x}_w, \bar{x}_\ell) &:= (\alpha\bar{x}_\ell - \beta(N-2)\bar{x}_\ell - \beta\bar{x}_w + b) \end{aligned} \tag{15}$$

Thresholding of the noise $\eta(t)$ by some activation threshold $\theta_{\text{act}}$ yields another effectively positive contribution, and to obtain the steady state ([2]) means we thus compute the effective means $\tilde{\mu}_{w/\ell}$ of a thresholded Gaussian noise with mean $\mu_w$ or $\mu_\ell$, respectively, and variance $\sigma_\eta^2$.

Due to linearity, these means $\tilde{\mu}_{w/\ell}$ must equal the average stationary activities $\bar{x}_{w/\ell}$. Simple integration yields

$$\begin{aligned} \tilde{\mu}_{w/\ell}(x_w, x_\ell) &= \int_{\theta_{\text{act}}}^{\infty} \frac{x}{\sqrt{2\pi}\sigma_\eta}\, e^{-\left(x-\mu(x_w,x_\ell)\right)^2/2\sigma_\eta^2}\, dx \\ &= \frac{\sigma_\eta}{\sqrt{2\pi}}\, e^{-\left(\theta_{\text{act}}-\mu(x_w,x_\ell)\right)^2/2\sigma_\eta^2} + \frac{\mu(x_w,x_\ell)}{2}\, \text{erfc}\left[\frac{(\theta_{\text{act}} - \mu(x_w,x_\ell))}{\sqrt{2}\sigma_\eta}\right] \end{aligned} \tag{16}$$

where erfc[.] denotes the complementary error function, and $\theta_{\text{act}} = 0$ if not explicitly stated otherwise.

Numerical fixed-point computation reveals the existence of two self-consistent solutions, depending on the noise amplitude $\sigma_\eta$: for no or small noise levels there is one branch of solutions $(\bar{x}_w, \bar{x}_\ell)$ with $\bar{x}_w \sim \frac{b_1}{1-\alpha} \gg \bar{x}_\ell \sim 0$, the *WTA-branch*, while for larger noise amplitudes another branch of solutions comes into existence, the *non-WTA-branch*, characterized by finite asymptotic rates of the same order, $\bar{x}_w \gtrsim \bar{x}_\ell$, see Figure S3a,b.

**Impact of input distribution:** The steady state model Equation (16) can be extended to arbitrary input distributions $b_i$, only limited by numerical stability of the fixed-point calculation: e.g., if all neurons but the top two are uniformly distributed, i.e., $b_i \sim \text{U}[0, 1-\Delta), i \geq 3$, the $(N-2)$ neurons with input drive $< (1-\Delta)$ are less likely to cross the activation threshold (driven by noise fluctuations) than for $b_{i\leq 2} \equiv 1-\Delta$. Thus, the whole WTA system tolerates an increased noise amplitude $\sigma_\eta$. For the same reason noisy WTA dynamics with uniformly distributed bulk drive are also faster than quasi-2d inputs: fluctuation driven inhibition can never be suppressed by the winner, so it will always have a slowing pull, but less so when fewer neurons contribute it in the first place.

**Hysteresis:** If the onset of the non-WTA-branch as a function of $\sigma_\eta$ is much earlier than the breakdown of the WTA-branch, hysteresis can be observed: if dynamics starts close to

[2] This approximation assumes $\tau_\eta \ll \tau$, such that the noise fluctuates on much faster time scales than the activities $x_i(t)$. If $\tau_\eta \gtrsim 0.01\tau$ the noise fluctuations become correlated with the fluctuations of $x_i(t)$. In this case a more involved self-consistent framework that takes into account the history of the dynamics needs to be employed.

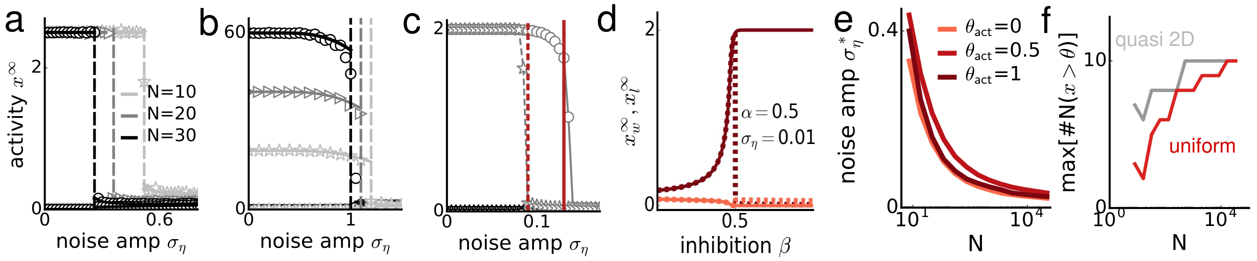

Figure S3: **Breakdown of WTA as a function of $\sigma_\eta$ and rescue by nonlinear inhibition** (a) For strong inhibition (here, $\beta = 1$, $\alpha = 0.6$) and increasing noise amplitude $\sigma_\eta$ the WTA solution (WTA branch, left of dashed line indicating the respective $\sigma_\eta^*$) with one neuron at $x_1^\infty = b_1/(1-\alpha)$ and all others at $x_{i\geq 2} = 0$ loses stability and breaks down, while a non-WTA solution (non-WTA branch) with all neurons asymptotically active becomes stable (right of dashed line). For increasing $N$ this transition moves to smaller $\sigma_\eta$ (light to dark gray). Symbols denote simulation results, lines theory. Other parameters: $b_1 = 1$, $b_{i\geq 2} = 0.9$, $\tau_\eta = 0.005\tau$. (b) Same for weak inhibition (note that the activity of loser neurons stays at finite levels in the WTA regime due to the positive bias of thresholded input noise). (c) If the WTA branch loses stability later than the non-WTA-branch gains stability, hysteresis can be observed: if initial conditions are close to the WTA solution the transition to non-WTA occurs later (solid red line) than if initial conditions are homogeneous across neurons (dashed red line). Parameters: $N = 100$, $\tau_\eta = 0.005\tau$. (d) Self-consistent solutions as a function of $\beta$ for fixed $\sigma_\eta = 0.01$, $N = 32$, $\Delta = 0.01$ and $\alpha = 0.5$. (e) Critical noise amplitude for different activation thresholds as obtained from the self-consistent theory. WTA dynamics gains a bit of stability for intermediate thresholds, but decreases again for high thresholds (higher threshold corresponds to darker hue). Parameters: $\alpha = 0.5$, $\beta = 0.6$, $\Delta = 0.05$, $\tau_\eta = 0.005\tau$, $\theta_{\mathrm{act}} = 0$. (f) Average maximal number of neurons providing nonlinear inhibition (i.e., neurons $j$ such that $[x_j(t) - \theta] > 0$) at the same time in steps of $\tau$: gray lines for quasi-2D drive, red lines for uniform drive. Parameters: $\alpha = 0.5$, $\beta = 0.6$, $\sigma_\eta = 0.2$, $\theta = 0.2$; averaged over 10 trials.

the WTA initial condition $x_{\mathrm{WTA}}^\infty$, it will stay stable to larger noise amplitude than if it starts close to the non-WTA initial condition $x_{\mathrm{nonWTA}}^\infty$, see Figure S3c.

**Dynamical constraints on $\alpha, \beta$ for WTA with noisy inputs:** For deterministic WTA networks $\alpha < 1$ is required for stability, while $\alpha + \beta > 1$ is needed for existence of WTA [8]. By means of the mean field theory we find that these constraints are still the relevant ones for noisy WTA with linear inhibition. We checked various combinations of $(\alpha, \beta)$-tuples, including those with $\alpha + \beta < 1$ (linearly stable) and $\alpha > 1$ and found no differences on the necessary conditions, such as, e.g., an extended noise-stabilized regime for WTA, compared to the deterministic dynamics. Figure S3d summarizes the general picture that for $\alpha + \beta > 1$ a WTA-state can occur for $\sigma_\eta < \sigma_\eta^*$ (solid lines), while for $\alpha + \beta < 1$ only the non-WTA-branches (dashed) exist.

**Dependence of stability on activation threshold $\theta_{\mathrm{act}}$** We observed that for strong linear inhibition stability to noise strongly decreases with network size $N$ because of residual asymptotic noise-driven inhibition, see Figure 2f in the main manuscript. One solution to overcome this problem could be increasing the activation threshold $\theta_{\text{act}}$, such that weakly driven neurons are well below threshold on average and thus cease to contribute interfering inhibition. Analysis of the self-consistent solution reveals that there is some gain in stability for intermediate $\theta_{\text{act}}$, but it is mainly a shift, not a general stabilization in the large-$N$ limit, see Figure S3e. Also, if the threshold is too big, stability is lost again, since not only losing neurons will fail to receive sufficient drive, but also a potential winner. Thus it is necessary to add a second inhibitory threshold rather than simply increasing the activation threshold.

**Source of noise robustness of nWTA with nonlinear inhibition** The reason for the increased robustness of nWTA to noise in the inputs is that only a few neurons effectively contribute inhibition at all in the presence of an inhibitory threshold, and that the maximum of this number of co-actively inhibiting neurons at any time during the computation depends on $N$ only weakly, and not very much on the distribution of $b_i$, see Figure S3f. Note that this number of active inhibiting neurons does depend on $\alpha, \beta$ and $\theta$, since these parameters determine how many neurons with $x(t) \geq \theta$ suffice to suppress all others.

#### S2.6 Speed-accuracy trade-off in noisy nWTA-networks with strong nonlinear inhibition

**Dependence of speed-accuracy trade-off on $\alpha$ and $\theta$** There are essentially three knobs to modulate speed and accuracy of nWTA networks: we can vary mutual inhibition $\beta$, self-excitation $\alpha$ or the inhibitory threshold $\theta$. In the main manuscript we vary $\beta$ for fixed $\alpha, \theta$. Here we consider the effect of each possible manipulation.

As with changing $\beta$, changing $\alpha$ allows the network a broad tradeoff between speed and accuracy, Figure S4a–c. The key difference of varying $\alpha$ instead of $\beta$ is that a variation of $\alpha$ modulates the asymptotic winner-activity: large $\alpha$ corresponds to higher final winner-activity. This explains why for increasing $\alpha$ the speed eventually decreases again, see Figure S4a–c. Thus for very high $\alpha$ the WTA system enters a regime of long computing time (for $\alpha \to 1$ $T_{\text{WTA}}$ diverges) at low accuracy (strong self-feedback can boost small random fluctuations in any neuron's activity and push it above the inhibitory threshold to suppress all other neurons).

The main effect of increasing the inhibitory threshold, on the other hand, is a reduction in accuracy, while speed and stability of WTA increase, see Figure S4a-f. Higher inhibitory threshold corresponds to a longer period of isolated integration driven by $b_i$ and reinforced by $\alpha$. The lack of mutual inhibition speeds this initial phase up. In the most extreme case, the neuron that reaches the inhibitory threshold $\theta$ first could then prohibit every other neuron from reaching $\theta$ themselves. This extreme case is hence akin to a race-to-threshold scenario. At high $\theta \geq 0.4$, accuracy is lower for higher numbers of competitors $N$ and hardly depends on $\alpha$, see Figure S4c. At the same time, the lack of competition stabilizes WTA, since the effect of noise-driven inhibition is minimized, Figure S4f. For the other extreme of low $\theta$ the linear inhibition WTA is recovered, with its low robustness to noise, i.e., high rate of WTA failure, see Figure S4a,d. At intermediate $\theta$ the width of the speed-accuracy trade-off is broadest, see Figure S4b versus a,c.

Similar results are obtained when varying $\beta$ and $\theta$, see Figure S4g–l.

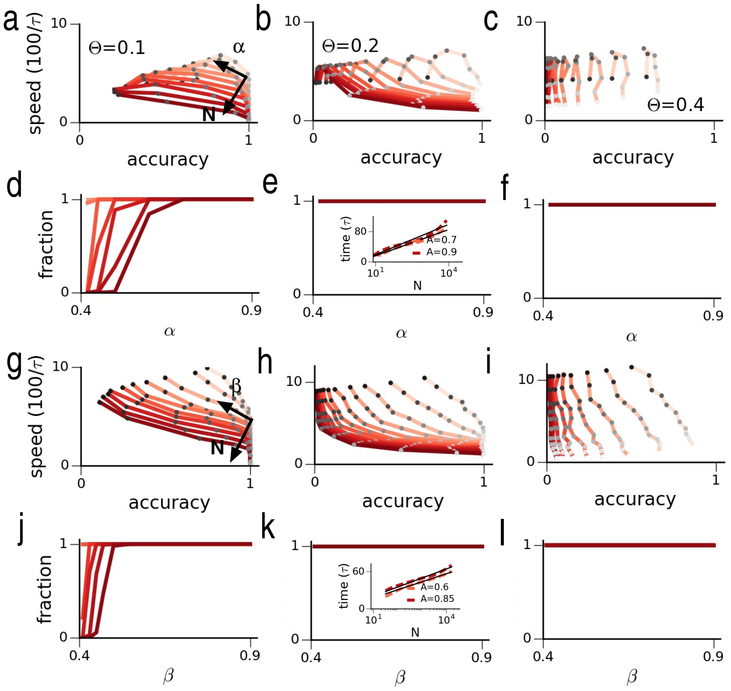

Figure S4: **Speed-accuracy trade-off as function of $\alpha$ and $\theta$** (a-c) Speed versus accuracy for increasing $N$ (light to dark red) and increasing self-excitation $\alpha = \{0.42, 0.45, 0.5, 0.6, 0.7, 0.8, 0.9\}$ (light to dark gray circles) for increasing inhibitory threshold $\theta = \{0.1, 0.2, 0.4\}$ (left to right). (d-f) Corresponding fraction of successful WTA. Other parameters: $\beta = 0.6$, $\sigma_\eta = 0.17$, $\tau_\eta = 0.05$, $\Delta = 0.05$. Note that only successful WTA cases were included into the speed-accuracy curves in (a-c). Inset in (e) shows the $N$-scaling of $T_{\text{WTA}}$ for two different fixed accuracies $A = 0.7, 0.8$ compared to a logarithm (dashed: simulations, solid: fit $\sim \log(N)$). (g–l) Same as (a–f) but varying $\beta$ along the curves at lower noise amplitude $\sigma_\eta = 0.15$ for fixed $\alpha = 0.6$. Inset in (k) shows logarithmic time scaling at fixed accuracies $A = \{0.6, 0.85\}$.

#### S2.7 Improvement of nWTA accuracy by noise

In the main text we show that the network exhibits a non-monotonic dependence of accuracy on noise level: except at very small values of $N$, the accuracy minimum occurs not at highest noise level, but at intermediate values (Figure 3c). Figures S5a-d show how $T_{\text{WTA}}$ (a,b) and accuracy (c,d) depend on $\Delta$ and $\sigma_\eta$, showing that for decreasing signal-to-noise ratio (larger $\sigma_\eta$ or also smaller $\Delta$) $T_{\text{WTA}}$ can increase to alleviate or even reverse accuracy deterioration.

Functionally the reason for the non-monotonicity of accuracy as a function of noise is as follows: For very low noise, dynamics are practically deterministic (leftmost panel of Figure S5e). At intermediate, but lower noise levels there are fewer neurons that contribute fluctuation driven inhibition, thus more neurons reach significant activation, compete and can win, decreasing accuracy and subtly increasing decision time (see Figure S5e, panel 2). At higher noise level more neurons contribute inhibition, which first (depending on parameters) slightly decreases (Figure S5e, panel 3), then strongly increases decision time (Figure S5e, panel 4), effectively increasing integration time and allowing the biased neuron to become

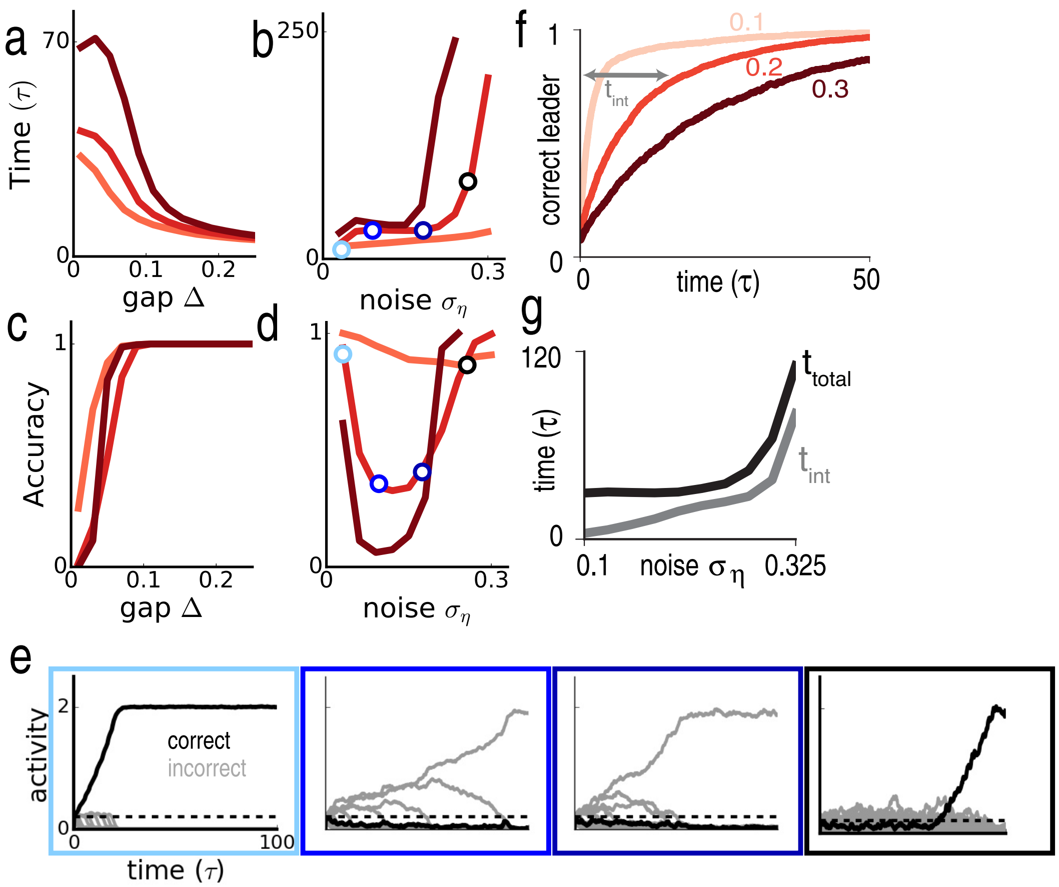

Figure S5: **Improvement by noise in nWTA with strong inhibition** (a,b) Mean $T_{\rm WTA}$ and (c,d) mean accuracy as functions of $\Delta$ and $\sigma_\eta$ for $N = \{8, 256, 16384\}$ (light to dark red). Network integrates for longer as task gets harder (smaller $\Delta$ or larger $\sigma_\eta$) and accuracy is non-monotonic with $\sigma_\eta$. (e) Example dynamical traces at increasing noise amplitude (from left to right, indicated by colored boxes and circles). Black solid trace: biased neuron; gray: all other neurons; dashed black line: inhibitory threshold $\theta$. (f) Fraction of trials on which the neuron with highest firing rate at time $t$ (shown on x-axis) is the eventual winner, as a function of $t$. Curves show three different values of noise ($\sigma_\eta$). Integration time is measured as the time by which 80% of trials have converged, meaning that the eventual winner is the same as the leader at that time. (g) Integration time and total decision time shown as function of noise. Parameters: throughout $\beta = 0.6$, $\tau_\eta = 0.05\tau$, $b_1 = 1, b_{i\geq 2} = 1 - \Delta$, $\theta = 0.2$. For panels (a)-(e): $\alpha = 0.5$, $\Delta = 0.075$ when held fixed, $\sigma_\eta = 0.15$ when held fixed. For panels (f), (g): $\alpha = 0.45$, $\Delta = 0.05$, $N = 20$.

the correct winner.

Figures S5f,g demonstrate that the improvement in performance with increasing noise amplitude is indeed due to a concurrent increase in the effective time over which the network is able to integrate input. The time taken by the network computation can be broken up into the time the network is responsive to input and an additional time during which the network converges to its final state, but is either unresponsive or weakly-responsive to input.

We probe the duration of integration by looking at the fraction of trials on which the neuron with the highest firing rate at time $t$ is the eventual winner (Figure S5f). For small $t$ this fraction should be close to chance, and once the computation is finished this fraction should be one. A simple measure of the integration window is the time at which the eventual winner can be predicted with 80% accuracy (i.e., 80% of trials have finished integrating): both this integration window and the total decision time increase with noise, Figure S5g, causing an initial increase in accuracy before the higher noise leads to lower accuracy. Thus, the most accurate trials are those where noise is large enough to extend the integration window but not large enough to cause a breakdown in accuracy.

### S3 WTA in small networks with noise: Multi-alternative forced-choice decision making

#### S3.1 Linear versus nonlinear inhibition

In Figures 4 and 5 we show results for decision-making between a small number of options with nonlinear inhibition. Results are similar for linear inhibition, see Figure S6, with some minor differences: The location of the parameters that produce best reward rate at $T_0 = 30\tau$, $\tau = 10\,\text{ms}$, depend more strongly on $N$ for nonlinear than linear inhibition, see Figure S6a,e. The reward rate scales slightly better for linear networks (Figure S6b,d,i), and accuracy at fixed time is higher (Figure S6c, upper panel), while $T_{\text{WTA}}$ at fixed, near-perfect accuracy is almost identical (Figure S6c, lower panel). Hick's law behavior for fixed parameters is less generic for linear inhibition than for nonlinear inhibition (see discussion below).

In Figure S6b we also include plots for the diff-AB strategy (gray lines), which considers the difference between the largest and second-largest integrated alternative, and terminates the decision when this crosses a threshold [6]. This strategy requires identifying the top two alternatives (out of $N$) at each moment, thus multiplying the time taken by $N$. However, if this additional time complexity is neglected and only the integration time of each option is considered, then diff-AB is optimal in the limit of vanishing error rate [6]. In Figure S6b we show the time taken by diff-AB when ignoring this additional time. For more discussion of diff-AB see S1.4.

##### S3.1.1 Optimal $(\alpha, \beta)$-tuples for small noisy WTA networks with linear and nonlinear inhibition

Figures S6e,f show how the best all-over values $(\alpha, \beta)^{\text{opt}}$ (lines) relate to the individually best (stars and circles) concretely. For $T_0 = 0$ best all-over and individual are very close for both linear and nonlinear inhibition networks, see Figure S6e. For $T_0 = 300$ the agreement is still good for linear inhibition, but values deviate for nonlinear inhibition: best $\alpha$ moves from small values to intermediate ones with increasing $N$, while best $\beta$ decreases from high to intermediate. Meanwhile, the best all-over values are almost averaging the individually bests, see Figure S6f. The reason for this difference in linear and nonlinear inhibition networks lies in the fact that for increasing $T_0$ speed becomes less relevant than accuracy in optimizing reward rate. Inspection of accuracy as a function of $\alpha, \beta$ (see Figure S6a for $N = 6$) reveals

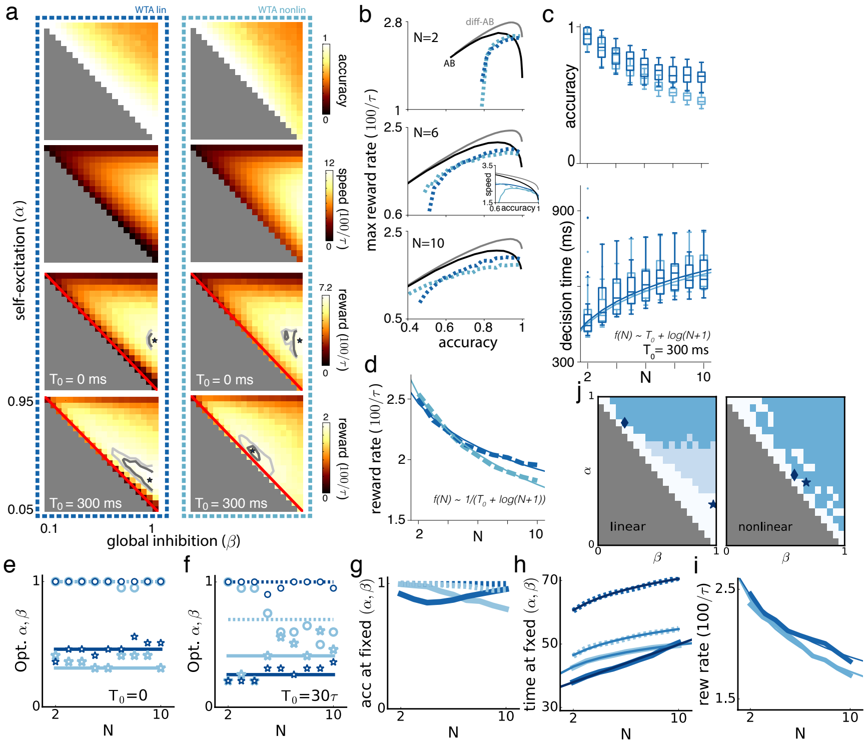

Figure S6: **Linear and nonlinear inhibition for multi-alternative forced choice decision-making** (a) Accuracy, speed and reward rate as a function of $\beta$ and $\alpha$ [$N = 6$; 4000 trials per tuple]; left column is identical to heatmaps in Figure 5, main text. Dark blue lines denote linear, light nonlinear inhibition networks in all panels. (b) Reward rate against accuracy for three different network sizes N = 2, 6, 10; dashed: WTA, solid: integration to threshold (AB and diff-AB). Inset in $N = 6$ panel shows speed-accuracy curves: solid shows varying $\alpha$, fixed $\beta$; dashed shows varying $\beta$, fixed $\alpha$. Note that dark blue solid curve lies mostly on top of dark blue dashed curve. (c) Upper panel: accuracy at fixed decision time $T_{\text{WTA}} = 90\,\text{ms}$. Lower panel: decision time at fixed accuracy $A = 0.99$ as a function of $N$; solid lines: fit of median values to $f(N) \sim \log(N+1)$. (d) Maximal reward rate as a function of $N$ at best individual parameters. (e) $\alpha$ (stars) and $\beta$ (circles) yielding maximal reward rate for each individual $N$. Solid and dashed lines indicate all-over best $(\alpha, \beta)^{\text{opt}}$ values which maximize reward rate across all $N$ [linear: $\alpha_{\text{opt}} = 0.46, \beta_{\text{opt}} = 1$; nonlinear: $\bar{\alpha} = 0.31, \bar{\beta} = 1$ at $T_0 = 0\,\text{ms}$]. Note that the two dashed lines lie on top of one another. (f) Same as (e) for $T_0 = 300\,\text{ms}$ with best parameters [linear: $\alpha_{\text{opt}} = 0.26, \beta_{\text{opt}} = 1$; nonlinear: $\bar{\alpha} = 0.41, \bar{\beta} = 0.7$]. (g) Accuracy at fixed $(\alpha, \beta)$: solid lines at $(\alpha, \beta)^{\text{opt}}$ (star in j), dashed lines at some fixed $(\alpha, \beta)$ that produces high accuracy across $N$ (diamond in j). (h) Decision time for same tuples as in (g) (thin solid: best fits either logarithmic or linear). (i) Reward rate at $(\alpha, \beta)^{\text{opt}}$ (thin solid: fit to $\sim 1/(T_0 + \log(N+1))$). (j) Heatmaps showing for which fixed $(\alpha, \beta)$-tuples $T_{\text{WTA}}$ of $N$ is best fit by Hick's law ($f(N) \sim T_0 + \log(N+1)$, dark blue squares), a linear function ($f(N) \sim a + bN$, light blue squares) or neither (white). Left panel: network with nonlinear inhibition, right panel linear inhibition. See text for details. Parameters: $\tau_\eta = 0.05\tau$, $\Delta = 0.05$, $\sigma_\eta = 0.2$, $b = (1, 1 - \Delta, \ldots, 1 - \Delta)^\top$.

that accuracy decreases for high $\beta$ and small $\alpha$ if inhibition is nonlinear, while it remains high for linear inhibition. So while the optimum is similar for both linear and nonlinear inhibition for $T_0 = 0$, it moves to more intermediate $\alpha, \beta$ for increasing $N$ for nonlinear inhibition. Accuracy in the nonlinear network decreases for large $\beta$ and small $\alpha$, because presence of an inhibitory threshold favors whoever crosses threshold first to become the winner, see also Figure S4.

##### S3.1.2 WTA-performance and Hicks law for fixed network parameters $\alpha, \beta$

Figure S6g–i show accuracy, $T_{\text{WTA}}$ and reward rate as a function of $N$ for fixed $(\alpha, \beta)$ tuples: Accuracy at $(\alpha, \beta)^{\text{opt}}$ is close to one for both networks (solid lines), with a slightly convex shape for linear, and monotonic decrease for nonlinear networks (Figure S6g); for the time dependence Hick's law is recovered, if inhibition is nonlinear (Figure S6h, solid light blue), while for linear networks the best fit is a linear increase of $T_{\text{WTA}}$ with $N$ (solid dark blue). Figure S6i shows that even though $T_{\text{WTA}}(N)$ for fixed $(\alpha, \beta)^{\text{opt}}$ are quite different for linear and nonlinear inhibition, reward rates of $N$ can be fit well by the inverse of Hick's law $f(N) \sim [T_0 + \log(N+1)]^{-1}$ (expected, if $T_{\text{WTA}}$ follows Hick's law at approximately constant accuracy) for both linear and nonlinear networks.

To see how generally a logarithmic dependence of $T_{\text{WTA}}$ on $N$ (as given by Hick's law) can be observed for any fixed admissible $(\alpha, \beta)$-tuple, we analyzed when $T_{\text{WTA}}(N)$ is either fit best by Hick's law $f(N) \sim T_0 + \log(N + 1)$ (Figure S6j, dark blue squares), a linear function $f(N) \sim N$ (light blue squares) or neither (white squares). Quality of fit was assessed by the mean squared error (MSE): white-blue squares indicate MSE$< 0.1 (< 0.05)$ for linear (nonlinear) inhibition networks; in those cases where the fit to either linear or logarithmic was deemed good, the fit with lowest MSE was chosen. For networks with nonlinear inhibition a good fit to Hick's law was obtained for most $(\alpha, \beta)$ tuples (Figure S6j, right panel), while for linear inhibition networks (left panel) that was the case only for larger $\alpha > 0.65$, while intermediate $\alpha$ and larger $\beta$ values gave rise to a linear $T_{\text{WTA}}(N)$. Stars indicate the $(\alpha, \beta)^{\text{opt}}$-tuples that gave best all-over performance with respect to reward rate optimization in Figure S6i, while diamonds mark tuples that give Hicks law at almost constant high accuracy (dashed lines in Figure S6g,h). These results show that a $\log(N)$-scaling of $T_{\text{WTA}}$ occurs for a wide variety of networks, especially for nonlinear inhibition, without changing any parameters with increasing $N$.

[4] Lutz Duembgen. Bounding standard gaussian tail probabilities. *arXiv preprint arXiv:1012.2063*, 2010.

[5] Wassily Hoeffding. Probability inequalities for sums of bounded random variables. *Journal of the American Statistical Association*, 58(301):13–30, 1963.

[6] Tyler McMillen and Philip Holmes. The dynamics of choice among multiple alternatives. *Journal of Mathematical Psychology*, 50(1):30–57, 2006.

[7] S Tajima, J Drugowitsch, and A Pouget. Optimal policy leads to irrational choice behavior under multiple alternatives. *Cosyne Abstracts 2017, Salt Lake City, UT, USA.*, 2017.

[8] X Xie, RHR Hahnloser, and SH Seung. Selectively grouping neurons in recurrent networks of lateral inhibition. *Neural Computation*, 14:2627–2646, 2002.
